# Deletion of a conserved Gata2 enhancer impairs haemogenic endothelium programming and adult haematopoiesis

**DOI:** 10.1101/516203

**Authors:** Tomasz Dobrzycki, Christopher B. Mahony, Monika Krecsmarik, Cansu Koyunlar, Rossella Rispoli, Joke Peulen-Zink, Kirsten Gussinklo, Bakhta Fedlaoui, Emma de Pater, Roger Patient, Rui Monteiro

## Abstract

Haematopoietic stem and progenitor cells (HSPCs) maintain the vertebrate blood system throughout life and their emergence from haemogenic endothelium (HE) is regulated by transcription factors such as Gata2. Here we deleted a conserved enhancer (i4 enhancer) driving pan-endothelial expression of *gata2a* and showed that Gata2a is required for HE programming by regulating expression of *runx1* and of the second zebrafish Gata2 orthologue, *gata2b*. By 5 days, homozygous *gata2a*^Δi4/Δi4^ larvae showed normal numbers of HSPCs, a recovery mediated by Notch signalling driving *gata2b* and *runx1* expression in HE. However, *gata2a*^Δi4/Δi4^ adults showed oedema, susceptibility to infections and marrow hypo-cellularity, consistent with bone marrow failure found in GATA2 deficiency syndromes. Thus, *gata2a* expression driven by the i4 enhancer is required for HE programming in embryos and maintenance of steady-state haematopoietic stem cell output in the adult. These enhancer mutants are a new paradigm to explore the pathophysiology of GATA2-related deficiencies *in vivo*.

## Introduction

Haematopoietic stem cells (HSCs) are the source of all blood produced throughout the lifetime of an organism. They are capable of self-renewal and differentiation into progenitor cells that generate specialised blood cell types. DNA-binding transcription factors are fundamental players in the inception of the haematopoietic system as it develops in the embryo, but also play a crucial role in maintaining homeostasis of the haematopoietic system in the adult organism. They are part of elaborate gene regulatory networks that coordinate differentiation, proliferation and survival of haematopoietic cells and ensure their levels are appropriate at all times throughout life. The only time during ontogenesis when HSCs are generated *de novo* is during embryonic development and misexpression of key transcription factors may lead to a failure to produce HSCs or, alternatively, to haematopoietic disorders and eventually leukaemia. Therefore, understanding how transcription factors drive the haematopoietic process provides opportunities for intervention when haematopoiesis is dysregulated.

The development of blood occurs in distinct waves: primitive, pro-definitive and definitive, each of them characterised by the generation of blood progenitors in a specific location and restricted in time, where the definitive wave produces multi-lineage self-renewing HSCs^1^. The specification of HSCs initiates in cells with arterial characteristics and proceeds through an endothelial intermediate, termed the haemogenic endothelium (HE)^2^. In zebrafish and other vertebrates, expression of *runx1* in the floor of the dorsal aorta defines the *bona fide* HE population ^3,4^. Haematopoietic stem and progenitor cells (HSPCs) emerge from the HE by endothelial-to-haematopoietic transition (EHT), both in zebrafish and in mice ^5-7^. They first arise at around 34 hours post fertilisation (hpf) from the HE in the ventral wall of the DA ^8^, the analogue of the mammalian AGM ^9^. After EHT, the HSCs enter the bloodstream through the posterior cardinal vein (PCV) ^8^ to colonise the caudal haematopoietic tissue (CHT), the zebrafish equivalent of the mammalian foetal liver ^10^. Afterwards the HSCs migrate again within the bloodstream to colonise the kidney marrow (WKM) and thymus ^8^, the final niche for HSCs, equivalent to the bone marrow in mammals ^1^.

Gata2 is a key haematopoietic transcription factor (TF) in development. In humans, *GATA2* haploinsufficiency leads to blood disorders, including MonoMAC syndrome (<u>Mono</u>cytopenia, <u>M</u>ycobacterium <u>a</u>vium <u>c</u>omplex) and myeloid dysplastic syndrome (MDS) ^11,12^. While its presentation is variable, MonoMAC syndrome patients always show cytopenias, ranging from mild to severe, and hypocellular bone marrow ^12,13^. These patients are susceptible to mycobacterial and viral infections and have a propensity to develop myelodysplastic syndrome (MDS) and Acute Myeloid Leukaemia (AML), with a 75% prevalence and relatively early onset at age 20 ^12^.

*Gata2* knockout mice are embryonic lethal and die by E10.5 ^14^. Conditional *Gata2* knockout under the control of the endothelial *VE-cad* promoter abolished the generation of intra-aortic clusters ^15^, suggesting that Gata2 is required for HSPC formation. In addition, conditional deletion of Gata2 mediated by a *Vav*-Cre transgene demonstrated that Gata2 is also required for the maintenance and survival of HSCs in the foetal liver, after HE specification and HSC emergence ^15^. Further studies in the mouse revealed a decrease in HSC numbers in *Gata2* heterozygous mutants, but also a dose-dependency of adult HSCs on Gata2 ^16^.

*Gata2* expression in the endothelium is regulated by an intronic enhancer element termed the +9.5 enhancer ^17,18^. Deletion of this enhancer results in the loss of HSPC emergence from HE, leading to lethality by E14 ^18^. The same element is also mutated in 10% of all the MonoMAC syndrome patients ^11^.

Because of a partial genome duplication during the evolution of teleost fish, numerous zebrafish genes exist in the form of two paralogues, including *gata2* ^19^. This provides an opportunity to separately identify the temporally distinct contributions made by each Gata2 orthologue. *Gata2a* and *gata2b* are only 57% identical and are thought to have undergone evolutionary sub-functionalisation from the ancestral vertebrate *Gata2* gene ^20,21^. *Gata2b* is expressed in HE in the floor of the dorsal aorta from 18hpf and is required for *runx1* expression in HE ^20^. In addition, lineage tracing experiments showed that *gata2b*-expressing HE cells gave rise to HSCs in the adult ^20^. Similar to the mouse Gata2, *gata2b* expression depends on Notch signalling and is a *bona fide* marker of HE, currently regarded as the functional ‘haematopoietic homologue’ of Gata2 in zebrafish^20^. By contrast, *gata2a* is expressed in all endothelial cells and in the developing central nervous system^20,22^. Homozygous *gata2a*^*um27*^ mutants showed arteriovenous shunts in the dorsal aorta at 48hpf ^23^. However, *gata2a* is expressed at 11hpf in the haemangioblast population in the posterior lateral mesoderm (PLM) that gives rise to the arterial endothelial cells in the trunk^24^, well before *gata2b* is expressed in HE. This suggests that *gata2a* might play a role in endothelial and HE programming and thus help to elucidate an earlier role for Gata2 in HSC development.

Here we show that the *gata2a* locus contains a conserved enhancer in its 4^th^ intron, corresponding to the described +9.5Kb enhancer in the mouse Gata2 locus^17,18^, a feature that was not found in the *gata2b* locus. Using CRISPR/Cas9 genome editing, we demonstrated that this region, termed the i4 enhancer, is required for endothelial-specific *gata2a* expression. Analysis of homozygous mutants (*gata2a*^Δi4/Δi4^ mutants) showed decreased expression of the HE-specific genes *runx1* and *gata2b*. Thus, endothelial expression of *gata2a*, regulated by the i4 enhancer, is required for *gata2b* and *runx1* expression in the HE. Strikingly, their expression recovers and by 48hpf, the expression of haematopoietic markers in *gata2a*^Δi4/Δi4^ mutants is indistinguishable from wild type siblings. We have demonstrated that this recovery is mediated by an independent input from Notch signalling, sufficient to recover *runx1* expression in HE and thus HSPC emergence by 48hpf. We conclude that *runx1* and *gata2b* are regulated by two different inputs, one Notch-independent input from Gata2a and a second from the Notch pathway, acting as a fail-safe mechanism for the initial specification of HSPCs in the absence of the input by Gata2a. Despite the early rescue, *gata2a*^Δi4/Δi4^ adults showed increased susceptibility to infections, oedema, a hypocellular kidney marrow (WKM) and neutropenia, a phenotype resembling key features of GATA2 deficiency syndromes in humans. We conclude that Gata2a is required to maintain steady-state haematopoietic output from adult HSPCs and this function requires the activity of the i4 enhancer.

## Results

### A region of open chromatin in intron 4 of the zebrafish *gata2a* locus is specific to endothelium and highly conserved

Because Gata2 genes are duplicated in zebrafish, we set out to unpick the different roles Gata2a and Gata2b play during HSC generation and homeostasis by identifying their regulatory regions. Analysis of sequence conservation revealed that one region within the fourth intron of the zebrafish *gata2a* locus was conserved in vertebrates, including mouse and human (Fig. 1a-c). This region, which we termed ‘i4 enhancer’, corresponds to the endothelial +9.5 Gata2 enhancer identified previously in the mouse ^17,18^ and human^25^. Notably, the *gata2b* locus did not show broad conservation in non-coding regions (Supplementary Fig. 1a).

**Figure 1.**
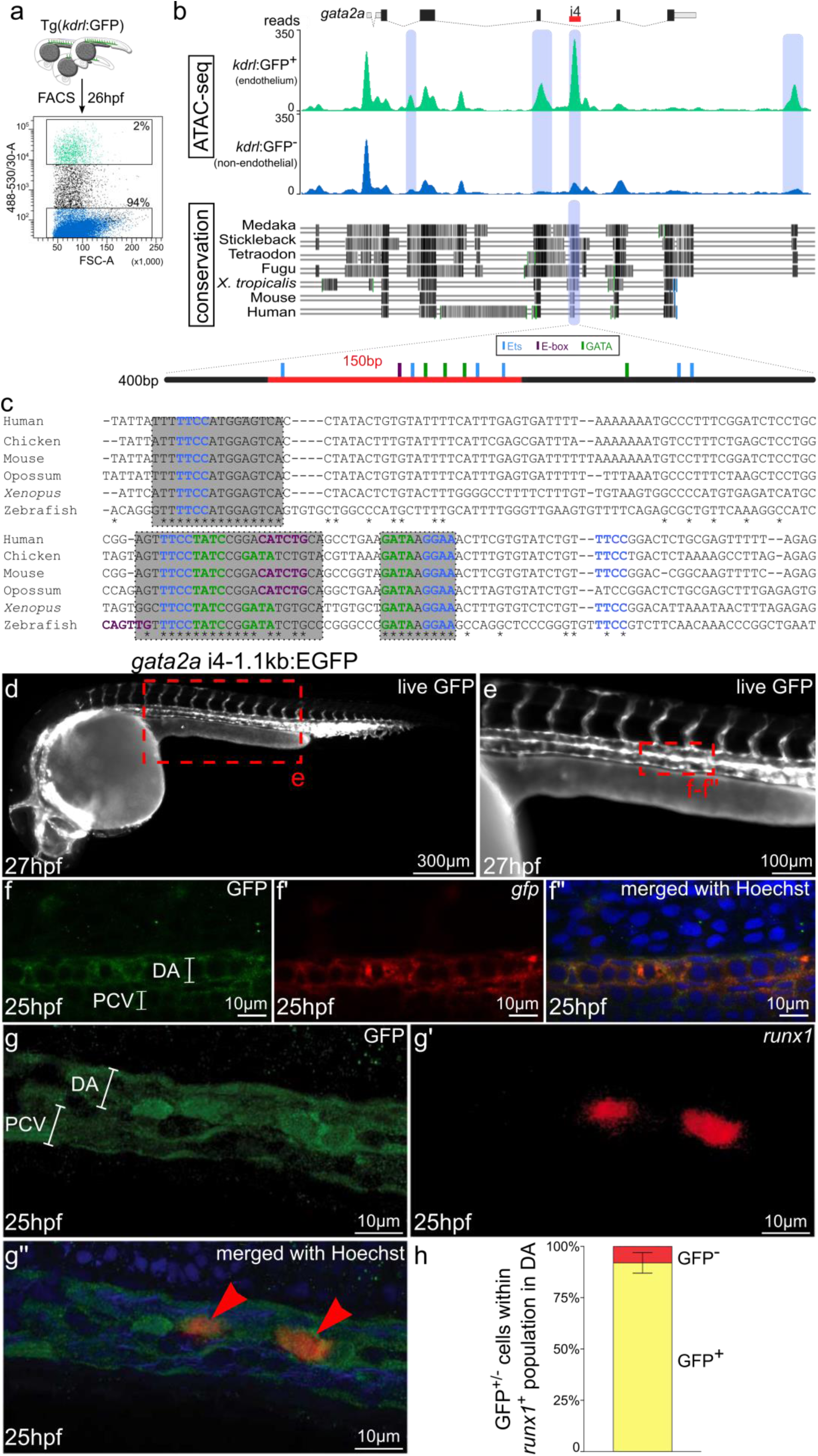
The i4 enhancer in the *gata2a* locus is conserved and drives pan-endothelial expression of a GFP reporter in zebrafish. (a) *Kdrl*:GFP^+^ (green) and *kdrl*:GFP^-^ (blue) cells were FACS-sorted from 26hpf embryos and used for preparation of ATAC-seq libraries. (b) The image of the mapped reads represents stacked means of two biological ATAC-seq replicates. Differential peak analysis identified four chromatin regions (blue shading) in the locus of *gata2a* that are significantly more open in the *kdrl*:GFP^+^ population (*p*<0.0001). A region in the fourth intron (termed i4 enhancer) is conserved throughout vertebrates. Black and grey shading denotes regions of high conservation between the species analysed (c) The highly conserved 150bp region (red) contains putative transcription factor binding sites, mapped computationally. Light blue: Ets binding sites; purple: E-box binding sites; green: GATA binding sites; asterisks: conserved residues. (d) Widefield fluorescent image of a live Tg(*gata2a*-i4-1.1kb:GFP) zebrafish embryo at 27hpf showing GFP fluorescence in the endothelial cells and in the heart (endocardium). (e) Higher magnification image of the trunk of the embryo from panel d. (f-f’’) Confocal images of a trunk fragment of a Tg(*gata2a*-i4-1.1kb:GFP) embryo immunostained with anti-GFP antibody (f) and probed for *gfp* mRNA (f’) at 25hpf. (f”) Merged images from panels f-f’ with Hoechst nuclear staining in blue, showing complete overlap of GFP protein and mRNA. (g-g’’) Confocal images of the dorsal aorta (DA) and posterior cardinal vein (PCV) of a Tg(*gata2a*-i4-1.1kb:GFP) embryo immunostained with anti-GFP antibody (g) and probed for *runx1* mRNA (g’) at 25hpf. See panel e for approximate position within the embryo. (g”) Merged images from panels g-g’, also showing Hoechst nuclear staining in blue. (h) Counting of the *runx1*^+^ cells represented in panels g’-g” in 25 embryos shows that >90% of *runx1*^+^ cells are also GFP^+^. N=3. Error bars: ±SD. See also Supplementary Figure 1.

To investigate whether the i4 element was a potentially active enhancer, we first performed ATAC-seq ^26^ to identify open chromatin regions in endothelial cells (ECs) in zebrafish. We used a Tg(*kdrl*:GFP) transgenic line that expresses GFP in all endothelium ^27^ and isolated the higher GFP-expressing ECs (*kdrl:GFP*^*high*^, termed *kdrl:GFP*^*+*^ for simplicity) as this fraction was enriched for endothelial markers compared to the *kdrl:GFP*^*low*^ fraction (Supplementary Fig. 1b,c). Principal Component Analysis on the ATAC-seq data from 26hpf *kdrl*:GFP^+^ cells (n=2) and *kdrl*:GFP^-^ cells (n=4) revealed strong differences between the open chromatin regions in the two cell populations, further supported by a correlation analysis (Supplementary Fig. 1d-f). 78,026 peaks were found in common between replicates of the ATACseq in *kdrl*:GFP^+^ cells (Supplementary Fig. 1g). 44,025 peaks were differentially expressed between the *kdrl:GFP*^*+*^ and *kdrl*:GFP^-^ fractions (Supplementary Fig. 1h). An analysis of known motifs present in the *kdrl:GFP*^*+*^ population revealed an enrichment for the ETS motif (Supplementary Fig. 1i). ETS factors are essential regulators of gene expression in endothelium ^28^. In addition, we performed gene ontology (GO) term analysis on the peaks showing >3-fold enrichment or depletion in ECs (Supplementary Fig. 1j-l). As expected, non-ECs showed a broad range of GO terms whereas EC-enriched peaks were associated with terms like angiogenesis or blood vessel development (Supplementary Fig. 1k,l).

Differential peak analysis in the *gata2a* locus identified four differentially open sites within a 20kb genomic region (Fig. 1b), including one peak in intron 4 corresponding to the predicted i4 enhancer. It contained a core 150bp-long element that included several binding motifs for the Gata, E-box and Ets transcription factor families (Fig. 1b). Although the positioning of the E-box site relative to the adjacent GATA site differs in zebrafish and mammals (Fig. 1b,c), the necessary spacer distance of ∼9bp between the two sites ^29^ was conserved. Thus, this site may be a target for TF complexes containing an E-box-binding factor and a GATA family TF.

Thus, the intronic enhancer (i4) identified in the zebrafish *gata2a* locus is accessible to transposase in endothelial cells and contains highly conserved binding sites for key haematopoietic transcription factors, suggesting that genetic regulation of *gata2a* expression in zebrafish HE is a conserved feature of vertebrate *gata2* genes.

### The *gata2a*-i4 enhancer drives GFP expression in the endothelium, including the haemogenic endothelium

To investigate the activity of the *gata2a*-i4 enhancer *in vivo*, the conserved genomic 150bp region (Fig. 1b,c), together with flanking ±500bp (*gata2a*-i4-1.1kb:GFP) or ±150bp (*gata2a*-i4-450bp:GFP) was cloned into a Tol2-based reporter E1b:GFP construct ^30^ and used to generate stable transgenic lines (Supplementary Fig. 2). The earliest activity of the enhancer was observed at the 14-somite stage (14ss), when *gfp* mRNA was detected in the PLM (Supplementary Fig. 2a,b). After 22hpf, the reporter signal was pan-endothelial (Fig. 1d-e, Supplementary Fig. 2c-i). Around 27hpf, higher intensities of GFP fluorescence and corresponding higher levels of *gfp* mRNA were visible in the floor of the DA (Fig. 1d-e, Supplementary Fig. 2e-h). While the GFP protein was still visible in the vasculature around 3dpf, it was likely carried over from earlier stages, since the *gfp* mRNA was not detectable anymore (Supplementary Fig. 2i,j). We focussed our subsequent analysis on the *gata2a*-i4-1.1kb:GFP transgenics as they showed stronger expression of the transgene (Supplementary Fig. 2k). At 25hpf, the expression of GFP protein and *gfp* mRNA overlapped completely in the endothelial cells of the DA (Fig. 1f-f’’). Overall, these data confirm that the i4 enhancer is active *in vivo* in endothelial cells at the correct time to regulate definitive haematopoiesis. The endothelial activity of the corresponding +9.5 enhancer was also observed in mouse embryos^17^, indicating functional conservation of the *gata2a*-i4 enhancer across vertebrates.

**Figure 2.**
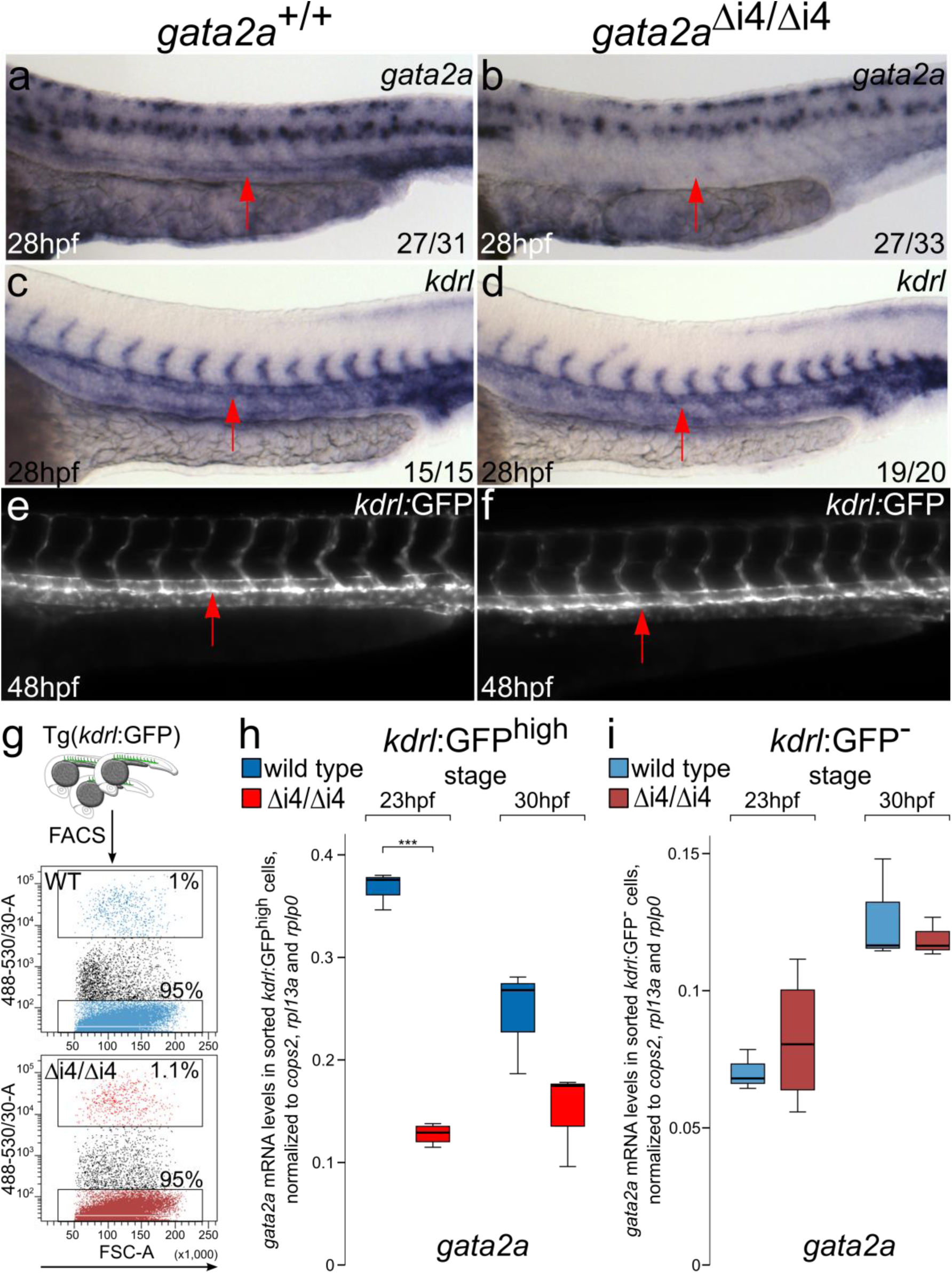
Deletion of the i4 enhancer in *gata2a*^Δi4/Δi4^ mutants leads to reduced levels of *gata2a* mRNA in the endothelium. (a-b) A significant majority of *gata2a*^Δi4/Δi4^ mutants have reduced levels of *gata2a* mRNA in the dorsal aorta (arrows) at 28hpf, compared to wild type siblings, as detected with *in situ* hybridization. (*X*^2^=10.720, d.f.=1, *p*<0.01) \*\**p*<0.01. The expression in the neural tube appears unaffected. (c-d) *In situ* hybridization for the endothelial marker *kdrl* at 28hpf reveals no difference between *gata2a*^Δi4/Δi4^ mutants and wild type siblings. The dorsal aorta (arrows) appears unaffected. (e-f) Live images of the trunks of 48hpf Tg(*kdrl*:GFP) and Tg(*kdrl*:GFP); *gata2a*^Δi4/Δi4^ embryos show normal vascular morphology in the mutants. The endothelium of the dorsal aorta (arrows) appears normal in the *gata2a*^Δi4/Δi4^ embryos. (g) *Kdrl*:GFP^high^ and *kdrl*:GFP^-^ cells were sorted from non-mutant (WT, blue) and *gata2a*^Δi4/Δi4^ (red) embryos carrying the Tg(*kdrl*:GFP) transgene. (h-i) qRT-PCR on RNA isolated from the sorted *kdrl*:GFP^high^ or *kdrl*:GFP^-^ cells (panel g) shows decreased levels of *gata2a* mRNA in the endothelium of *gata2a*^Δi4/Δi4^ mutants at 23hpf (*t*=20.026,d.f.=5, p<0.001) compared to wild type. At 30hpf this difference is not statistically significant (*t*=2.146, d.f.=4, p=0.098). There is no difference in *gata2a* mRNA levels in non-endothelial cells between wild type and *gata2a*^Δi4/Δi4^ mutants (23hpf: t=0.69, d.f.=5, p>0.5; 30hpf: t=0.618, d.f.=4, p>0.5). N=4 for *gata2a*^Δi4/Δi4^ at 23hpf, N=3 for other samples. Note different scales of expression levels. \*\*\**p*<0.001. See also Supplementary Figure 2.

To further characterise the enhancer activity *in vivo*, Tg(*gata2a*-i4-1.1kb:GFP) embryos were stained for *gata2a* mRNA and for GFP protein (Supplementary Fig. 2k-o). We found a large overlap between *gata2a*^+^ and GFP^+^ cells at 30hpf in the DA, with a small proportion of GFP^+^ cells that did not express *gata2a* mRNA (<5%, Supplementary Fig. 2o). This could suggest that some cells require activity of other endothelial enhancers to trigger transcription of *gata2a* or that *gfp* mRNA has a longer half-life than *gata2a* mRNA. Importantly, the GFP signal was absent in *gata2a*-expressing neural cells (Supplementary Fig. 2l-n), indicating that the i4 enhancer is specifically active in (haemogenic) endothelial cells.

Next we examined the expression of the HE marker *runx1* ^3^ in *gata2a*-i4-1.1kb:GFP embryos at 25hpf. At this stage, over 90% of *runx1*^+^ cells were GFP^+^ (Fig. 1g-h). We conclude that the GFP expression under the *gata2a*-i4 enhancer marks the majority of the HE population.

### Deletion of the *gata2a*-i4 enhancer results in decreased expression of *gata2a* in the embryonic endothelium without affecting endothelial development

To investigate whether endothelial-specific expression of *gata2a* is required for definitive haematopoiesis, we deleted the conserved *gata2a*-i4 enhancer using CRISPR/Cas9 genome editing ^31^. We generated a deletion mutant lacking 231bp of the i4 enhancer (Supplementary Fig. 3a-c) and named it *gata2a*^Δi4/Δi4^. Homozygous *gata2a*^Δi4/Δi4^ mutants showed decreased levels of *gata2a* expression in endothelial cells when compared to wild type embryos (Fig. 2a,b). By contrast, *gata2a* expression in the neural tube appeared unaffected in the *gata2a*^Δi4/Δi4^ mutants (Fig. 2a,b). At 28hpf, expression of the pan-endothelial marker *kdrl* was indistinguishable between wild type and *gata2a*^Δi4/Δi4^ mutants (Fig. 2c,d). To verify these results, we crossed homozygous *gata2a*^Δi4/Δi4^ mutants to Tg(*kdrl*:GFP) transgenics and analysed vascular morphology. *Gata2a*^Δi4/Δi4^ embryos showed no gross vascular abnormalities at 48hpf as assessed by the expression of the Tg(*kdrl*:GFP) transgene (Fig. 2e,f).

**Figure 3.**
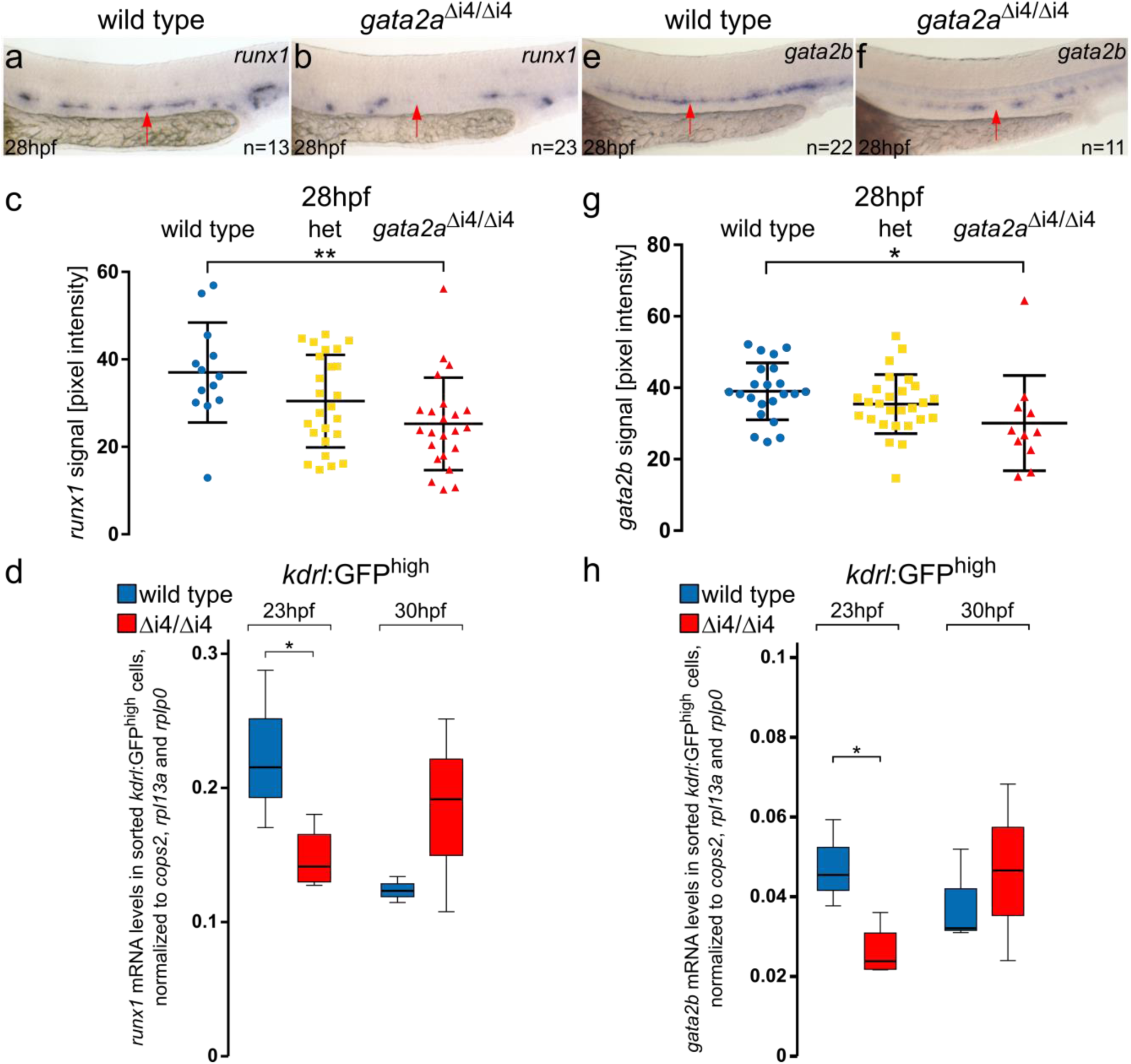
Loss of gata2a expression in the endothelium of *gata2a*^Δi4/Δi4^ mutants leads to decreased levels of *runx1* and *gata2b* in the HE. (a-b) *In situ* hybridization for *runx1* expression in the HE (arrows) of wild type and *gata2a*^Δi4/Δi4^ embryos at 28hpf. (c) Quantification of the *runx1 in situ* hybridization signal from wild type (blue), heterozygous *gata2a*^+/Δi4^ (het, yellow) and *gata2a*^Δi4/Δi4^ (red) siblings at 28hpf shows significant decrease in *runx1* pixel intensity in the DA in the homozygous mutants compared to wild type (*μ*_wt_=34.8, *μ*_mut_=25.3; *F*=4.956, d.f.=2, 58; ANOVA),\*\**p*<0.01. n=14, wild type; n=25, het; n=23, *gata2a*^Δi4/Δi4^. Error bars: mean±SD. (d) qRT-PCR on RNA isolated from the sorted *kdrl*:GFP^+^ cells shows decreased levels of *runx1* mRNA in the endothelium of *gata2a*^Δi4/Δi4^ mutants at 23hpf (*t*=2.585, d.f.=5, *p*<0.05) but not at 30hpf (t=1.326, d.f.=4, p>0.2), compared to wild type. N=4 for *gata2a*^Δi4/Δi4^ at 23hpf, N=3 for other samples. Note different scales of expression levels. \**p*<0.05. (e-f) *Gata2b* expression in the HE (arrows) of wild type and *gata2a*^Δi4/Δi4^ embryos at 28hpf. (g) Quantification of the *gata2b* mRNA signal, detected by *in situ* hybridization, from wild type (blue), heterozygous *gata2a*^+/Δi4^ (het; yellow) and *gata2a*^Δi4/Δi4^ (red) siblings at 28hpf shows significant decrease in *gata2b* pixel intensity in the DA in the homozygous mutants compared to wild type (*μ*_wt_=39, *μ*_mut_=30.1; *F*=5.05, d.f.=2, 54; ANOVA), \**p*<0.05. n=22, wild type; n=24, het; n=11, *gata2a*^Δi4/Δi4^. Error bars: mean±SD. (h) qRT-PCR in sorted *kdrl*:GFP^+^ cells showed decreased levels of *gata2b* mRNA in the endothelium of *gata2a*^Δi4/Δi4^ mutants at 23hpf (*t*=3.334, d.f.=5, *p*<0.05) but not at 30hpf (t=0.373, d.f.=4, p>0.7), compared to wild type. N=4 for *gata2a*^Δi4/Δi4^ at 23hpf, N=3 for other samples. \**p*<0.05. See also Supplementary Figures 3 and 4.

Next, we isolated endothelial cells from Tg(*kdrl*:GFP) and Tg(*kdrl*:GFP); *gata2a*^Δi4/Δi4^ embryos by FACS (Fig. 2g) at 23hpf and 30hpf and confirmed by qRT-PCR that the endothelial markers *kdrl, dld* and *dll4* were unaffected in *gata2a*^Δi4/Δi4^ embryos (Supplementary Fig. 3d-f). The arterial marker *efnb2a* ^32^ was decreased at 23hpf in *gata2a*^Δi4/Δi4^ mutants but recovered by 30hpf (Supplementary Fig. 3g). In addition, *gata2a* was significantly decreased in the *kdrl*:GFP^+^ ECs in 23hpf *gata2a*^Δi4/Δi4^ embryos compared to wild types (Fig. 2h). At 30hpf this decrease was not statistically significant. This was likely due to a decrease in expression of *gata2a* that appears to occur in wild type ECs during development, whereas *gata2a* expression in mutants remained low (Fig. 2h). Importantly, there was no difference in *gata2a* expression in the non-endothelial population (*kdrl*:GFP^-^ cells) between wild type and *gata2a*^Δi4/Δi4^ mutants at either 23hpf or 30hpf (Fig. 2i). Altogether, these data suggest that genomic deletion of the *gata2a*-i4 enhancer is sufficient to reduce expression of *gata2a* specifically in endothelium.

### Deletion of the *gata2a*-i4 enhancer results in reduced expression of *runx1* and *gata2b* at early stages of HE specification

To investigate a potential role of *gata2a* in HSC development, we compared the expression of *runx1*, the key marker of HE in zebrafish ^3^, in wild type and *gata2a*^Δi4/Δi4^ embryos. Quantitative *in situ* hybridization (ISH) analysis ^33^ showed that *runx1* expression was decreased in *gata2a*^Δi4/Δi4^ embryos at 24hpf (Supplementary Fig. 4a-c) and 28hpf compared to wild type siblings (Fig. 3a-c). Further analysis in *kdrl*:GFP^+^ ECs showed that this decrease in *runx1* expression was already detectable at 23hpf in *gata2a*^Δi4/Δi4^ mutants (Fig. 3d), at the onset of its expression in HE ^34^. Thus, deletion of the *gata2a*-i4 enhancer results in impaired *runx1* expression in the early stages of HE programming. This correlates well with decreased *runx1* expression levels in +9.5^-/-^ mouse AGM explants ^18^, further supporting the critical evolutionary role of the intronic enhancer of *Gata2* in HSC specification.

**Figure 4.**
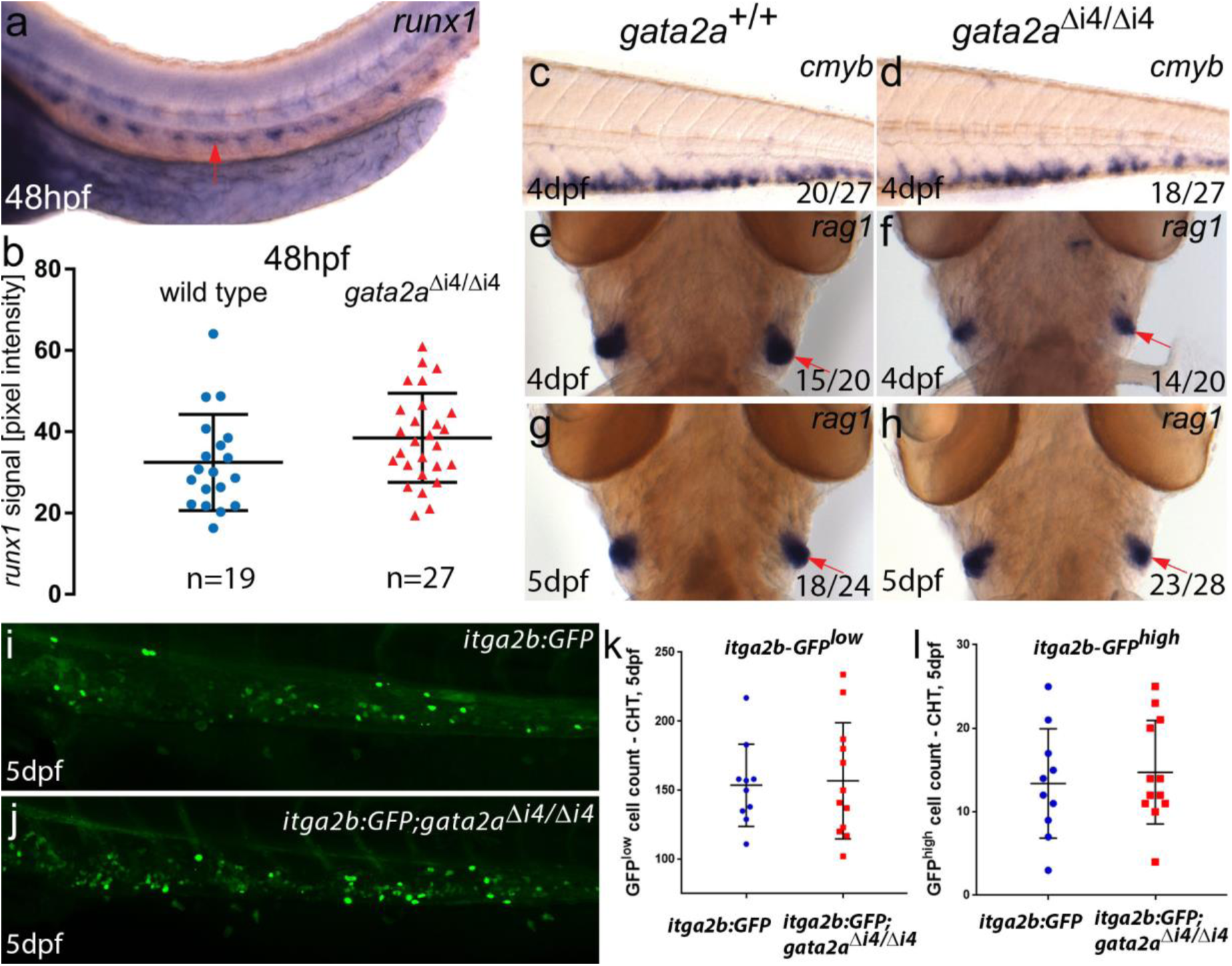
*Gata2a* ^Δi4/Δi4^ mutants display a recovery of the initial haematopoietic defects from 48hpf. (a) Representative image of *runx1* expression in the trunk of a wild type embryo at 48hpf showing *runx1* mRNA in the dorsal aorta (arrow). (b) Quantification of the *runx1 in situ* hybridization signal in wild type (blue) and *gata2a*^Δi4/Δi4^ mutants (red) siblings at 48hpf. There is no significant difference in *runx1* pixel intensity in the DA between the homozygous mutants and wild type (*μ*_wt_=33.1, *μ*_mut_=37.5, *t*=1.410, d.f.=44, p=0.17. n=19, wild type; n=27, *gata2a*^Δi4/Δi4^). Error bars: mean±SD. (c-d) *In situ* hybridization for *cmyb* in the CHT. We detected no difference in expression between wild type and *gata2a*^Δi4/Δi4^ siblings at 4dpf. (e-h) *In situ* hybridization (ventral image) for *rag1* in the thymii, showing a slight decrease (relative to wild type) in *rag1* (red arrows) in approximately half of the homozygous mutant embryos at 4dpf. This effect is absent at 5dpf. (i-j) Maximum projections of *itga2b*:GFP transgenic embryos in the CHT at 5dpf in (i) wild type and (j) *gata2a*^Δi4/Δi4^ siblings. (k) HSPC (*itga2b*:GFP^low^) counts in the CHT of wild type (n=10) and *gata2a*^Δi4/Δi4^ mutants (n=12) at 5dpf. No difference was detected between genotypes (μ_wt_=153.5; μ_mut_=145.5; p=0.98, Mann-Whitney test). (l) Thrombocyte (*itga2b*:GFP^high^) counts in the CHT of wild type (n=10) and *gata2a*^Δi4/Δi4^ mutants (n=12) at 5dpf. No difference was detected between genotypes (μ_wt_=13; μ_mut_=13; p=0.71, Mann-Whitney test). The scatter plots show the median±SD.

Next, we tested whether Gata2a could act upstream of *gata2b* by measuring *gata2b* expression in *gata2a*^Δi4/Δi4^ embryos. Quantitation of the *ISH* signal showed that *gata2b* expression was decreased in *gata2a*^Δi4/Δi4^ embryos compared to wild type siblings at 26hpf (Supplementary Fig. 4d) and 28hpf (Fig. 3e-g), but recovered to wild type levels by 30hpf (Supplementary Fig. 4e). Accordingly, *kdrl*:GFP^+^; *gata2a*^Δi4/Δi4^ cells express significantly lower levels of *gata2b* mRNA than the wild type *kdrl*:GFP^+^ endothelial population in 23hpf embryos, but not at 30hpf (Fig. 3h). These data suggest that endothelial expression of *gata2a* is required upstream of *gata2b* and *runx1* for the proper specification of HE, uncovering a previously unrecognized role for Gata2a in definitive haematopoiesis.

### Expression of late markers of embryonic HSC activity is unaffected in *gata2a*^Δi4/Δi4^ mutants

The qPCR analysis in sorted *kdrl*:GFP^+^; *gata2a*^Δi4/Δi4^ and wild type *kdrl*:GFP^+^ ECs (Fig. 3d) already suggested a recovery of of *runx1* expression from 30hpf. Of note, the *kdrl*:GFP^+^ population likely includes the *kdrl*^+^, *runx1*-expressing EMPs located in the caudal region ^35^. This region was not included in the quantification of *ISH* but cannot be separated by sorting for *kdrl*:GFP^+^ and could thus explain the discrepancy between image quantification and qRT-PCR. To further characterize the haematopoietic phenotype in the *gata2a*^Δi4/Δi4^ mutants, we tested whether expression of markers of haematopoietic activity in the embryo was affected from 48hpf onwards (Fig. 4).

At 48hpf, the expression of *runx1* in the DA showed no significant difference between *gata2a*^Δi4/Δi4^ mutants and wild type controls (Fig. 4a,b). These data suggest that the decrease of *runx1* expression at early stages of HE programming in *gata2a*^Δi4/Δi4^ mutants is transient and recovers by 2dpf. Indeed, analysis of the HSPC marker *cmyb* ^10^ in the CHT at 4dpf showed no differences between *gata2a*^Δi4/Δi4^ and wild type larvae (Fig. 4c,d). Expression of the T-cell progenitor marker *rag1* in the thymus ^36^ showed that around half of the *gata2a*^Δi4/Δi4^ larvae had reduced *rag1* expression at 4dpf compared to wild type (Fig. 4e,f). This effect was absent at 5dpf (Fig. 4g,h), suggesting that HSPC activity was normal in *gata2a*^Δi4/Δi4^ mutants from 4dpf onwards. Next, we crossed the *gata2a*^Δi4/Δi4^ mutants to Tg(*itga2b*:GFP) transgenics, where *itga2b*-GFP^high^ and *itga2b*-GFP^low^ cells in the CHT mark thrombocytes and HSPCs, respectively ^8,37^. Our analysis revealed no difference in *itga2b*-GFP^low^ HSPC or *itga2b*-GFP^high^ thrombocyte numbers in the CHT region at 5dpf between wild type and *gata2a*^Δi4/Δi4^ mutants (Fig.4i-l). Taken together, our data suggest that endothelial *gata2a* expression mediated by the i4 enhancer is required for the initial expression of *gata2b* and *runx1* in the HE but largely dispensable after 2dpf.

### Haematopoietic recovery of *gata2a*^Δi4/Δi4^ mutants is mediated by the Notch-*gata2b* pathway

The recovery of *gata2b* expression by 30hpf (Fig. 3h, Supplementary Fig. 4e) coincides temporally with the observed decrease in *gata2a* in wild type endothelial cells (Fig. 2h). Thus, we reasoned that other regulators of *gata2b* might compensate for the lack of endothelial *gata2a* in *gata2a*^Δi4/Δi4^ mutants and thus lead to a recovery of the initial haematopoietic phenotype. Therefore, we investigated whether the loss of *gata2b* in *gata2a*^Δi4/Δi4^ background resulted in a more severe haematopoietic phenotype than observed in the *gata2a*^Δi4/Δi4^ mutants. For this, we injected *gata2a*^Δi4/Δi4^ and wild type controls with a sub-optimal amount (7.5ng) of a *gata2b* morpholino oligonucleotide (MO) ^20^. Quantitative ISH analysis confirmed that this amount of *gata2b* MO had no effect on *runx1* expression at 32hpf (Fig. 5a,b). As expected, *runx1* expression in *gata2a*^Δi4/Δi4^ embryos was significantly reduced compared to wild type siblings (Fig. 5a,b). *Gata2b* knockdown in *gata2a*^Δi4/Δi4^ embryos further reduced *runx1* expression (Fig. 5a,b). To test whether this stronger reduction of *runx1* at 32hpf affected later stages of embryonic haematopoiesis, we assessed *cmyb* expression in the CHT at 4dpf (Fig. 5c). We scored *cmyb* expression levels as ‘wild type’ or ‘reduced’ and found that the ‘reduced’ embryos were largely overrepresented in the *gata2a*^Δi4/Δi4^ mutants injected with the *gata2b* MO, compared to wild type fish and non-injected *gata2a*^Δi4/Δi4^ siblings (Fig. 5c).

**Figure 5.**
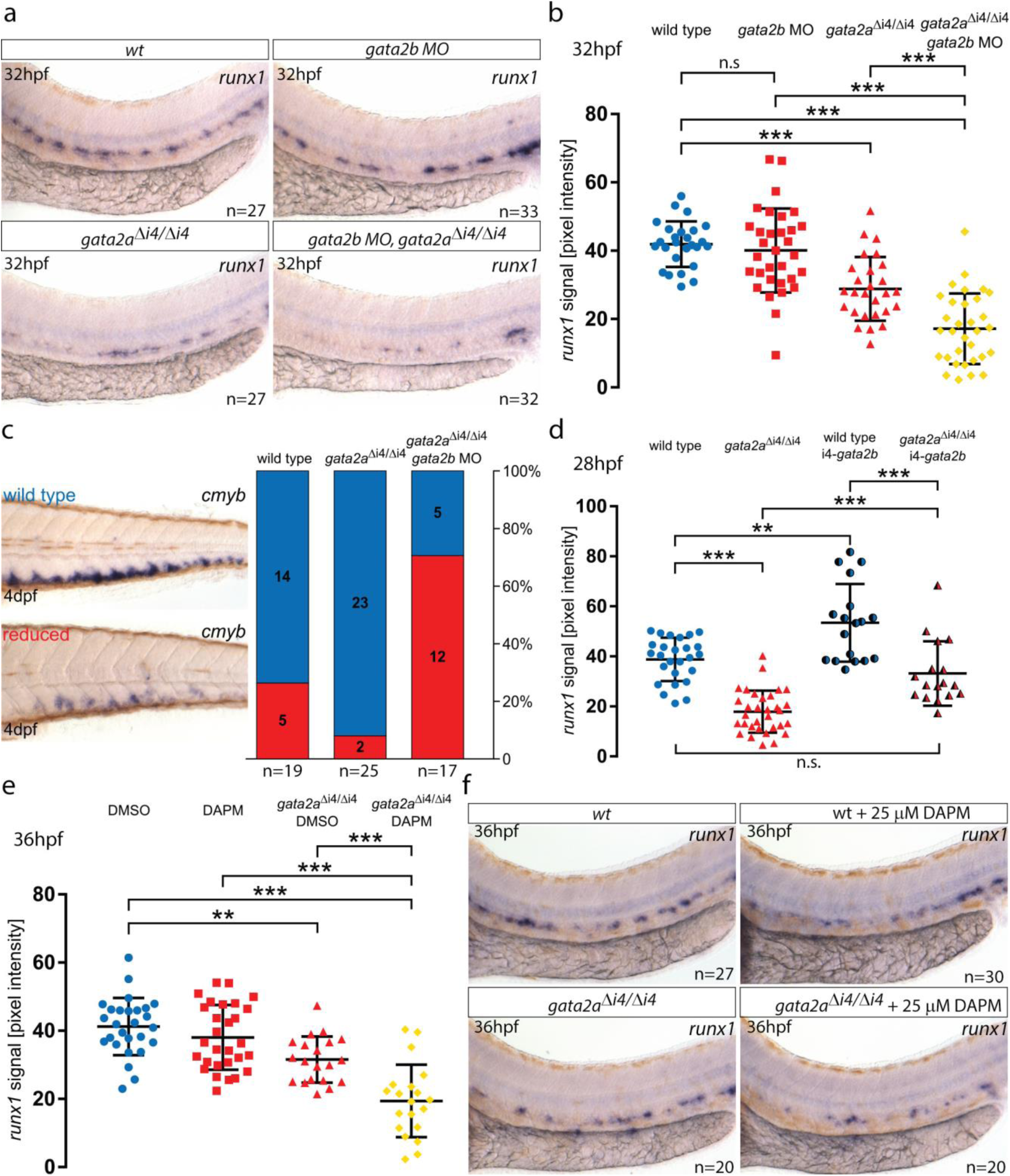
*Gata2b* and Notch signalling are sufficient to recover haematopoietic markers in *gata2a*^Δi4/Δi4^ mutants. (a) Expression of *runx1* in HE at 32hpf in wild type (wt), *gata2b MO*-injected (7.5ng) wt embryos, *gata2a*^Δi4/Δi4^ mutants and *gata2b MO*-injected (7.5ng) *gata2a*^Δi4/Δi4^ mutants. (b) Quantification of the *runx1 in situ* hybridization (ISH) signal in wt, *gata2b* morphants, *gata2a*^Δi4/Δi4^ mutants and *gata2a*^Δi4/Δi4^ mutants injected with *gata2b* MO. *runx1* expression is decreased in *gata2a*^Δi4/Δi4^ mutants (*μ*_wt_=41.9, *μ*_mut_=28.9; *F*=44.641, d.f.=3, 62.3; *p*<0.001). *Gata2b MO* knockdown significantly decreases *runx1* in the DA of *gata2a*^Δi4/Δi4^mutants (*μ*_mut_=28.9, *μ*_mut+MO_=17.2), but not wt embryos at 32hpf (*μ*_wt_=41.9, *μ*_MO_=40.1; p=0.89, p=0.89, Games-Howell post-hoc test, Welch’s ANOVA). n=27, wt; n=27, *gata2a*^Δi4/Δi4^; n=33, wt + *gata2b* MO; n=32, *gata2a*^Δi4/Δi4^ + *gata2b* MO. (c) Scoring *cmyb* expression at 4dpf in wt, *gata2a*^Δi4/Δi4^ mutants and *gata2a*^Δi4/Δi4^ mutant embryos injected with *gata2b* MO as wt (blue) or reduced (red). *Gata2b MO* knockdown (7.5ng) inhibits the haematopoietic recovery of *gata2a*^Δi4/Δi4^ mutants. (*X*^2^=18.784, d.f.=2,p<0.001). (d) Quantification of the *runx1* ISH signal, from 28hpf wt embryos (blue), *gata2a*^Δi4/Δi4^ mutants (red) and their siblings injected with a *gata2a*-i4-450bp:*gata2b* construct (shaded blue and red). Ectopic expression of *gata2b* increases *runx1* expression in the HE of wt embryos (*μ*_wt_=38.8, *μ*_wt+*gata2b*_=53.4; *p*<0.01) and rescues *runx1* expression in the DA of *gata2a*^Δi4/Δi4^ mutants to wt levels (*μ*_mut_=17.9, *μ*_mut+*gata2b*_=33.2; *p*<0.001; *μ*_wt_=38.8, *μ*_mut+*gata2b*_=33.2; p=0.31, Tukey HSD post-hoc test). n=25, wt; n=33, *gata2a*^Δi4/Δi4^; n=18, wt + *gata2a*-i4-450bp:*gata2b* construct; n=17, *gata2a*^Δi4/Δi4^ + *gata2a*-i4-450bp:*gata2b* construct. (e) Quantification of the *runx1* ISH signal at 36hpf in embryos treated with a suboptimal dose (25μM) of the Notch inhibitor DAPM. 25 μM DAPM showed no effect on *runx1* expression in wt compared to DMSO-treated embryos (*μ*_DMSO_=40.5, *μ*_DAPM_=38; p=0.735, Tukey HSD post-hoc test). DMSO-treated *gata2a*^Δi4/Δi4^ mutants show a decrease in *runx1* expression (*μ*_DMSO_=40.5, *μ*_mut+DMSO_=31.5; *F*=25.774, d.f.=3, 91; ANOVA). DAPM treatment significantly reduced *runx1* expression in the DA *gata2a*^Δi4/Δi4^ mutants (μ_mut+DMSO_=31.5, *μ*_mut+DAPM_=19.4). n=27, wt+ DMSO; n=20, *gata2a*^Δi4/Δi4^+ DMSO; n=30, wt+DAPM; n=20, *gata2a*^Δi4/Δi4^+ DAPM. (f) Representative images of the average *runx1* expression at 36hpf in wt and *gata2a*^Δi4/Δi4^ mutants treated with 25μM DAPM. Error bars: mean±SD. \*\**p*<0.01; \*\*\**p*<0.001. See also Supplementary Fig. 5.

To verify whether Gata2b is required for definitive haematopoiesis downstream of Gata2a, we generated a frameshift truncating mutant for Gata2b^38^ and incrossed *gata2a*^Δi4/+^; *gata2b*^*+/-*^ adults to investigate *cmyb* expression at 33hpf in their progeny. *Gata2b*^*-/-*^ mutants showed a more severe decrease in *cmyb* expression than *gata2a*^Δi4/Δi4^mutants (Supplementary Fig. 4f-i). Double *gata2b*^*-/-*^; *gata2a*^Δi4/Δi4^ mutants showed no further reduction in *cmyb* expression compared to *gata2b*^-/-^ mutants, suggesting that Gata2a was not sufficient to drive *cmyb* expression in HE in the absence of Gata2b (Supplementary Fig. 4f-i). Taken together, we conclude that Gata2b is regulated by Gata2a and is required for definitive haematopoiesis.

Next, we tested whether forced ectopic expression of *gata2b* was sufficient to speed up the haematopoietic recovery of *gata2a*^Δi4/Δi4^ embryos. Thus, we overexpressed *gata2b* under the control of the *gata2a*-i4-450bp enhancer in wild type and *gata2a*^Δi4/Δi4^ mutant embryos and measured *runx1* expression at 28hpf in the DA. *Gata2a*^Δi4/Δi4^ embryos showed a significant decrease in *runx1* expression in compared to wild type (Fig. 5d). Ectopic expression of *gata2b* under the *gata2a*-i4 enhancer significantly increased *runx1* expression in wild type and mutants (Fig. 5d). Importantly, it was sufficient to bring the *runx1* expression levels in the mutants up to the levels detected in uninjected wild type embryos (Fig. 5d), suggesting that *gata2b* alone was sufficient to drive *runx1* expression in the ventral wall of the DA and drive the haematopoietic recovery in *gata2a*^Δi4/Δi4^ mutants. Thus, *gata2b* can recover the definitive haematopoietic programme in the absence of endothelial *gata2a*.

Because the expression of *gata2b* is regulated by Notch signalling ^20^, we investigated whether inhibition of Notch would also prevent the haematopoietic recovery of *gata2a*^Δi4/Δi4^ embryos. For this, we used the Notch inhibitor DAPM ^39^, and titrated it down to a sub-optimal dose (25μM) that did not significantly affect *runx1* expression (Supplementary Fig. 5a). This dose induced a small but measurable decrease in *gata2b* expression in DAPM-treated embryos while higher doses had a more robust effect (Supplementary Fig. 5b). Next, we treated wild type and *gata2a*^Δi4/Δi4^ mutant embryos with DAPM and measured *runx1* expression in the DA at 36hpf (Fig. 5e,f). Suboptimal DAPM treatment did not affect *runx1* expression in wild type embryos (Fig. 5e,f), but *gata2a*^Δi4/Δi4^ mutants showed lower *runx1* levels and DAPM treatment further reduced *runx1* expression (Fig. 5e,f). By contrast, *gata2b* expression at 36hpf was unaffected by 25μM DAPM but strongly reduced by 100μM DAPM (Supplementary Fig. 5c). Taken together, these data show that Notch activity is sufficient to drive haematopoietic recovery in *gata2a*^Δi4/Δi4^ mutants. Consistent with these results, ectopic activation of Notch signalling in endothelium with a *fli1a*-NICD:GFP construct^40^ led to increased *runx1* and *gata2b* expression in wild type embryos at 30hpf (Supplementary Fig. 5d,e). When overexpressed in *gata2a*^Δi4/Δi4^ mutants, *fli1a*-NICD:GFP rescued *runx1* expression to near wild type levels at 26hpf (Supplementary Fig. 5f). Thus, we conclude that HE programming requires two independent inputs on *runx1* and *gata2b* expression; one from Gata2a, driven in ECs by the i4 enhancer, and the other from Notch signalling, necessary and sufficient to drive HE programming even in the absence of *gata2a*.

### Adult *gata2a*^Δi4/Δi4^ mutants show defects resembling GATA2 haploinsufficiency

To investigate whether Gata2a plays a role in adult haematopoiesis, we first asked whether the *gata2a*-i4-1.1kb:GFP reporter was active in haematopoietic cells in the adult. Whole kidney marrow (WKM) cells isolated from the transgenic fish showed that the i4 enhancer is active in haematopoietic cells previously defined by flow cytometry ^41^ as progenitors, lymphoid+HSPC (containing the HSPCs) and myeloid cells (Supplementary Fig. 6a-c). Accordingly, single cell transcriptional profiling showed higher levels of *gata2a* in HSPCs, progenitors, neutrophils and thrombocytes (Supplementary Fig. 6d-f) ^42,43^. Consistent with this notion, we observed a high incidence of infections and heart oedemas in *gata2a*^Δi4/Δi4^ adult fish, with over 25% suffering from one of these defects by 6 months of age, compared to <1% of wild type fish (Fig. 6a-c). The heart oedemas and the infections are suggestive of lymphatic defects and immune deficiency as observed in human patients bearing genetic GATA2 haploinsufficiency syndromes such as MonoMAC syndrome ^12^. Notably, around 10% of MonoMAC syndrome patients show mutations in the homologous enhancer region of *GATA2* ^11,13^.

**Figure 6.**
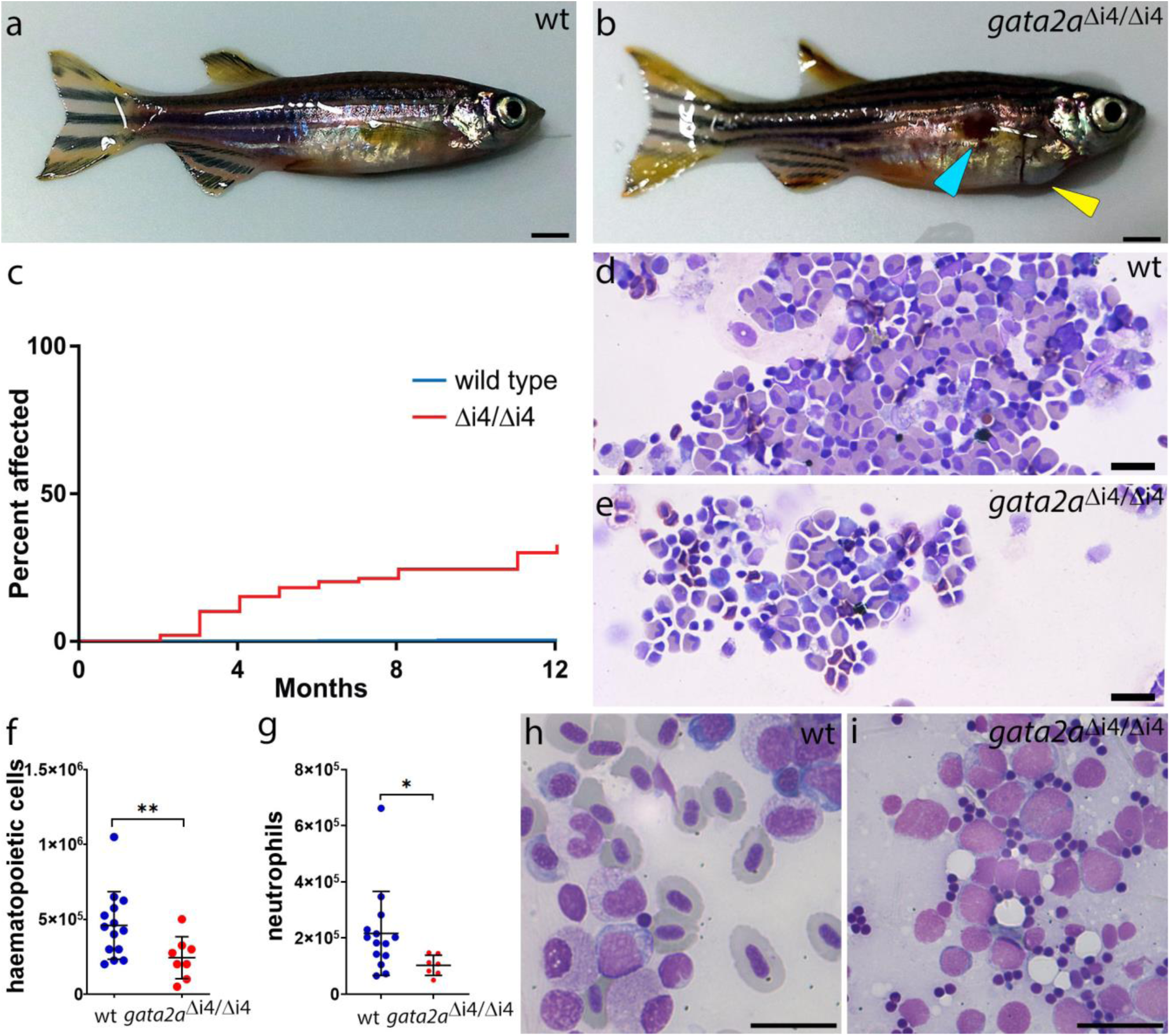
*Gata2a*^Δi4/Δi4^ mutants show cardiac oedema, hypocellularity and marrow failure. (a-b) General morphology of zebrafish adults: (a) wild type; (b) *gata2a*^Δi4/Δi4^ mutant showing skin infection (blue arrowhead) and pericardial oedema (yellow arrowhead). (c) Over 25% (n=29/108) of *gata2a*^Δi4/Δi4^ mutants (red) catch infections or suffer from heart oedemas by 6 months. Only around 65% (n=69/108) survive for more than 12 months without overt signs of infections. Fewer than 1% (n=2/500) of wild type fish (blue) exhibit such defects. The graph does not include deaths by other causes. (d-e) May-Grunwald/Wright-Giemsa staining in cytospins of haematopoietic cells isolated from the WKM of zebrafish adults: (d) wild type; (e) *gata2a*^Δi4/Δi4^ mutant. Note the decrease in cell numbers. (f) Cell counts of haematopoietic cells isolated from WKM of wild type (n=14) and *gata2a*^Δi4/Δi4^ mutants (n=8). The *gata2a*^Δi4/Δi4^ mutants show a ∼2-fold decrease in haematopoietic cell numbers in the WKM (μ_wt_=4.37×10^5^;μ_mut_=2.37×10^5^, *p*=0.0185, Mann-Whitney test). (g) Number of neutrophils isolated from WKM of wild type (n=14) and *gata2a*^Δi4/Δi4^ mutants (n=7). The *gata2a*^Δi4/Δi4^ mutants show a ∼2-fold decrease in neutrophil numbers in the WKM (μ_wt_=2.17×10^5^;μ_mut_=1.03×10^5^, *p*=0.0269, Mann-Whitney test). The scatter plots show the median cell number±SD. (h-i) Kidney smears from 9 months post-fertilization adult animals were assessed. (h) Wild type shows various stages of lineage differentiation. (i) WKM smear; 1 of 10 *gata2a*^Δi4/Δi4^ mutants showed the presence of excess blasts with very little erythroid differentiation (98% blasts, > 200 cells assessed). Scalebars: 2mm (a,b) and 10μm (d,e,h,i). See also Supplementary Fig.6.

Next, we counted the total number of haematopoietic cells in wild type and *gata2a*^Δi4/Δi4^ mutant WKM (Fig. 6d-f). To avoid any confounding effects in our analysis, we compared wild type to *gata2a*^Δi4/Δi4^ mutants without overt signs of infection. The *gata2a*^Δi4/Δi4^ mutants showed a ∼2-fold decrease in the total number of haematopoietic cells in the WKM (Fig. 6d-f). In addition, neutrophils were similarly reduced (Fig. 6g), another characteristic in common with MonoMAC syndrome patients ^13^. Lastly, kidney marrow smears of ten 9-month old *gata2a*^Δi4/Δi4^ mutants were assessed. One of the ten mutants showed an excess of immature myeloid blast cells in the WKM (>98%) and only minor erythrocyte differentiation (Fig. 6h,i). The presence of excess blasts is usually an indication of AML in humans. Together these data strongly suggest that the i4 enhancer is a critical driver of *gata2a* expression in adult haematopoietic cells. The enhancer deletion in *gata2a*^Δi4/Δi4^ mutants leads to a hypocellular WKM and neutropenia, strongly suggestive of marrow failure, a hallmark of disease progression in Gata2 deficiency syndromes.

## Discussion

### Endothelial expression of *gata2a* through a conserved intronic enhancer is required for HE programming

The sub-functionalisation of the Gata2 paralogues in zebrafish provided an opportunity to unpick the different roles of *Gata2* in the multi-step process of definitive haematopoiesis. Here we have investigated the conservation of the Gata2 +9.5 enhancer and identified a homologous region in intron 4 of the zebrafish *gata2a* locus (*gata2a*-i4) that is not present in the *gata2b* locus. The zebrafish *gata2a*-i4 enhancer, like the mouse enhancer ^17^, is sufficient to drive pan-endothelial expression of GFP and necessary for endothelial expression of *gata2a* (Fig. 1,2). We traced the activity of the i4 enhancer back to the PLM, the source of precursors of endothelium and HSCs ^9^. This degree of sequence and functional conservation of the i4 enhancer led us to hypothesize that Gata2a might play a role in definitive haematopoiesis. Indeed, homozygous deletion of the i4 enhancer (*gata2a*^Δi4/Δi4^) allowed us to uncover a previously unknown function of Gata2a in regulating the initial expression of *runx1* and *gata2b* in HE. Although *cmyb* expression in HE was decreased in *gata2a*^Δi4/Δi4^ mutants, it was more severely reduced in *gata2b*^*-/-*^ mutants, suggesting that Gata2b is more important for *cmyb* regulation than Gata2a. However, both Gata2 orthologues regulate gene expression in the HE before the first reported EHT events at 34hpf ^5^.

### Regulation of *gata2a* and *gata2b*

Gata2 binds to the +9.5 enhancer to maintain its own expression in endothelial and haematopoietic cells ^25,44^. In zebrafish, it is likely that Gata2a binds the GATA motifs in the i4 enhancer and loss of *gata2a* in the endothelium of *gata2a*^Δi4/Δi4^ mutants (Fig. 2) seems to support this view. Interestingly, we detected a small region in intron 4 of the *gata2b* locus that was not identified as a peak in our ATACseq experiment but is conserved in some fish species (Supplementary Fig. 1a) and thus could potentially represent a divergent gata2b intronic enhancer. We speculate that the positive autoregulation of Gata2 was likely retained by both *gata2* orthologues in zebrafish. In this case, Gata2a would bind to the *gata2b* intronic enhancer in HE only until enough Gata2b is present, at which point its intronic enhancer would ‘switch’ Gata2a for Gata2b to maintain *gata2b* expression. This ‘switch’, however, would replace one activator for another, rather than replacing an activating Gata factor (Gata2) for a repressive Gata factor as described previously for the Gata2/Gata1 switch in erythroid cells ^45^. Alternatively, the positive auto-regulation function might have been modified in HE so that Gata2a binds to both *gata2a* and *gata2b* intronic enhancers to regulate their expression. These possibilities remain to be investigated.

### Rescue of the haematopoietic defects in *gata2a*^Δi4/Δi4^ mutants

*The gata2a*^Δi4/Δi4^ mutants recovered from the early defects in HE programming and displayed normal expression levels of *cmyb* in the CHT at 4dpf and *rag1* in the thymus at 5dpf, used as indicators of the definitive haematopoietic programme ^10^. We hypothesized that this could be due to the presence of the two homologues of *Gata2* in zebrafish ^19^, despite Gata2a and Gata2b proteins being only 50% identical ^20^. Indeed, forced expression of *gata2b* under the *gata2a*-i4 enhancer rescued the expression of *runx1* in the *gata2a*^Δi4/Δi4^ mutants to wild type levels and sub-optimal depletion of *gata2b* in the *gata2a*^Δi4/Δi4^ mutants resulted in more severe reduction in *cmyb* expression in the CHT by 4dpf (Fig.4). In addition, we demonstrated that Notch signalling, a known regulator of *gata2b* expression ^20^, is sufficient to rescue the initial HE programming defect induced by deletion of the *gata2a*-i4 enhancer. We propose a model in which *gata2a* acts upstream of *runx1* and *gata2b* independently of Notch to initiate HE programming. The regulation of *gata2b* by Gata2a is transient, and the timing largely coincides with the natural decrease in endothelial expression of *gata2a* by 30hpf. After this stage, endothelial Notch signalling takes over the regulation of *runx1* and *gata2b* expression, acting as a fail-safe mechanism that buffers against fluctuations in the system caused by loss of one or more of the initial inputs (in this case, Gata2a).

In the mouse, +9.5^-/-^ enhancer mutants exhibit decreased haematopoietic output from cultured AGM explants ^18^ or foetal liver HSCs ^25^. In both cases the experiments were performed at E11.5, well after the HE had been established and given rise to HSCs ^4^. At E11.5 the numbers of haematopoietic cells in the AGM were roughly comparable to their wild type siblings ^18,25^, raising the possibility that an initial HE defect might have gone unnoticed in those studies. Thus, by analysing the contribution of two zebrafish Gata2 paralogues, we uncovered a previously unappreciated contribution by a Gata2 gene in the programming of HE prior to HSC specification.

### Loss of i4 enhancer activity leads to a phenotype resembling human GATA2 deficiency syndromes

Despite the apparent haematopoietic recovery, we observed a high incidence of infections and oedema in *gata2a*^Δi4/Δi4^ adults, and a striking decrease in the number of haematopoietic cells in the WKM. The decrease in haematopoietic cells in particular is reminiscent of the loss of proliferative potential of haematopoietic Gata2^+/-^ heterozygous cells in the mouse ^16,46^. This raises the possibility that in zebrafish the *gata2a* and *gata2b* paralogues may function as two Gata2 ‘alleles’ that together regulate the haematopoietic output of the WKM. This will be addressed by comparing the adult phenotypes of *gata2a*^Δi4/Δi4^ and *gata2b*^-/-^ mutants.

Taken together, our initial characterization of WKM shows that *gata2a*^Δi4/Δi4^ mutants present a phenotype consistent with Gata2 deficiency syndromes in humans brought about by GATA2 haploinsufficiency ^11,25^. Strikingly, about 10% of all MonoMAC patients show mutations in the conserved +9.5 enhancer ^11,13^, the corresponding regulatory element to the i4 enhancer. The i4 enhancer is active in the lymphoid+HSPC fraction that contains the HSC activity ^47^, in the progenitor cells and in the myeloid fraction that contains eosinophils, previously identified as expressing high levels of a *gata2a*-GFP BAC transgenic reporter ^41^. Thus, it is likely that *gata2a*^Δi4/Δi4^ adult fish show lineage-specific differentiation defects. Further characterization of the *gata2a*^Δi4/Δi4^ mutants will uncover which haematopoietic cells are most affected by the loss of i4 enhancer activity and how Gata2a regulates haematopoietic output, thus establishing a novel animal model for human diseases linked to *Gata2* haploinsufficiency.

## Methods

### Maintenance of zebrafish

Zebrafish (*Danio rerio*) were maintained in flowing system water at 28.5°C, conductance 450-550μS and pH 7.0±0.5 as described ^48^. Fish suffering from infections or heart oedemas were culled according to Schedule 1 of the Animals (Scientific Procedures) Act 1986. Eggs were collected by natural mating. Embryos were grown at 24-32°C in E3 medium with methylene blue and staged according to morphological features (Kimmel et al. 1995) corresponding to respective age in hours or days post fertilization (hpf or dpf, respectively). Published lines used in this work were wild type (wt^KCL^), Tg(−6.0*itga2b*:EGFP)^la2 37,49^ and Tg(*kdrl*:GFP)^s843 27^. All animal experiments were approved by the relevant University of Oxford, University of Birmingham and Erasmus University ethics committees.

### ATAC-seq

Tg(*kdrl*:GFP)^s843^ embryos were dissociated for FACS at 26-27hpf to collect *kdrl* ^+^ and *kdrl* ^-^ cell populations (40,000-50,000 cells each). They were processed for ATAC library preparation using optimised standard protocol ^26^. Briefly, after sorting into Hanks’ solution (1xHBSS, 0.25% BSA, 10mM HEPES pH8), the cells were spun down at 500g at 4°C, washed with ice-cold PBS and resuspended in 50μl cold Lysis Buffer (10mM Tris-HCl, 10mM NaCl, 3mM MgCl_2_, 0.1% IGEPAL, pH 7.4). The nuclei were pelleted for 10min. at 500g at 4°C and resuspended in the TD Buffer with Tn5 Transposase (Illumina), scaling the amounts of reagents accordingly to the number of sorted cells. The transposition reaction lasted 30min. at 37°C. The DNA was purified with PCR Purification MinElute Kit (QIAGEN). In parallel, transposase-untreated genomic DNA from *kdrl*^+^ cells was purified with the DNeasy^®^ Blood & Tissue Kit (QIAGEN). The samples were amplified with appropriate Customized Nextera primers ^26^ in NEBNext High-Fidelity 2x PCR Master Mix (NEB). The libraries were purified with PCR Purification MinElute Kit (QIAGEN) and Agencourt AMPure XP beads (Beckmann Coulter). The quality of each library was verified using D1000 ScreenTape System (Agilent). Four biological replicas of the libraries were quantified with the KAPA Library Quantification Kit for Illumina^®^ platforms (KAPA Biosystems). The libraries were pooled (including the Tn5-untreated control), diluted to 1ng/μl and sequenced using 75bp paired-end reads on Illumina HiSeq 4000 (Wellcome Trust Centre for Human Genetics, Oxford). Raw sequenced reads were checked for base qualities, trimmed where 20% of the bases were below quality score 20, and filtered to exclude adapters using *Trimmomatic* (Version 0.32) and mapped to Zv9 reference genome (comprising 14,612 genes) ^50^ using *BWA* with default parameters. The results were visualised using UCSC Genome Browser (http://genome-euro.ucsc.edu/) ^51^. The eight data sets were analysed with Principal Component Analysis (PCA) to identify outliers. Correlation among *kdrl:GFP* ^+^ and *kdrl:GFP* ^-^ samples was assessed with a tree map. The peaks were called for each sample using the Tn5-untreated control as input. We identified the common peaks between replicates and then used DiffBind (EdgeR method) to identify differential peaks between *kdrl:GFP* ^+^ and *kdrl:GFP* ^-^samples (Supplementary Table 2). The threshold for differential peaks was *p*<0.05.

### Generation of transgenic Tg(*gata2a-i4-1*.*1kb:GFP), Tg(gata2a-i4-450bp:GFP*) and mutant *gata2a*^**Δ**i4/**Δ**i4^ and *gata2b*^-/-^ zebrafish lines

Genomic regions containing the identified 150bp-long *gata2a*-i4 enhancer flanked by ±500bp (i4-1.1kb) or ±150bp (i4-450bp) were amplified from wild type zebrafish genomic DNA with NEB Phusion^®^ polymerase (see Supplementary Table 1 for primer sequences) and cloned upstream of E1b minimal promoter and GFP into a Tol2 recombination vector (Addgene plasmid #37845, ^52^) with Gateway^®^ cloning technology (Life Technologies™) following the manufacturer’s protocol. One-cell zebrafish embryos were injected with 1nl of an injection mix, containing 50pg *gata2a*-i4-E1b-GFP-Tol2 construct DNA + 30pg *tol2* transposase mRNA^30^. Transgenic founders (Tg(*gata2a*-i4-1.1kb:GFP) and (*gata2a*-i4-450bp:GFP)) were selected under a widefield fluorescent microscope and outbred to wt fish. Carriers of monoallelic insertions were detected by the Mendelian distribution of 50% fluorescent offspring coming from wt outcrosses. These transgenics were then inbred to homozygosity.

To generate the i4 deletion mutant, we identified potential sgRNA target sites flanking the 150bp conserved region within intron 4 of the *gata2a* locus (see Fig. 1A, Fig S3A). sgRNAs were designed with the CRISPR design tool (http://crispr.mit.edu/, see Supplementary Table 1 for sequences) and prepared as described ^31^. To reduce potential off-target effects of CRISPR/Cas9, we utilized the D10A ‘nickase’ version of Cas9 nuclease ^53,54^, together with two pairs of sgRNAs flanking the enhancer (Supplementary Table 1, Supplementary Fig. 3A-B). We isolated two mutant alleles with deletions of 215bp (Δ78-292) and 231bp (Δ73-303) (Supplementary Fig. 3B). Both deletions included the highly conserved E-box, Ets and GATA transcription factor binding sites (Supplementary Fig. 3B). The Δ73-303 allele was selected for further experiments and named Δi4. Adult zebrafish were viable and fertile as heterozygous (*gata2a*^Δi4/+^) or homozygous (*gata2a*^Δi4/Δi4^). To unambiguously genotype wild types, heterozygotes and homozygous mutants, we designed a strategy consisting of two PCR primer pairs (Supplementary Fig. 3A, C). One primer pair flanked the whole region, producing a 600bp wild type band and 369bp mutant band. In the second primer pair, one of the primers was designed to bind within the deleted region, only giving a 367bp band in the presence of the wild type allele (Supplementary Fig. 3C).

To generate the gata2b mutant we designed a CRISPR/Cas9 strategy for a frameshift truncating mutant in exon 3 deleting both zinc fingers. sgRNAs were designed as described above and guides were prepared according to Gagnon et al. ^55^ with minor adjustments. Guide RNAs were generated using the Agilent SureGuide gRNA Synthesis Kit, Cat# 5190-7706. Cas9 protein (IDT) and guide were allowed to form ribonucleoprotein structures (RNPs) at RT and injected in 1 cell stage oocytes. 8 embryos were selected at 24 hpf and lysed for DNA isolation. Heteroduplex PCR analysis was performed to test guide functionality and the other embryos from the injection were allowed to grow up. To aid future genotyping we selected mutants by screening F1 for a PCR detectable integration or deletion in exon 3. Sequence verification showed that founder 3 had a 28 nt integration resulting in a frameshift truncating mutation leading to 3 new STOP codons in the third exon. To get rid of additional mutations caused by potential off target effects, founder 3 was crossed to WT for at least 3 generations. All experiments were performed with offspring of founder 3.

### Fluorescence-activated cell sorting (FACS)

∼100 embryos at the required stage were collected in Low Binding^®^ SafeSeal^®^ Microcentrifuge Tubes (Sorenson) and pre-homogenized by pipetting up and down in 500μl Deyolking Buffer (116mM NaCl, 2.9mM KCl, 5mM HEPES, 1mM EDTA). They were spun down for 1min. at 500g and incubated for 15min. at 30°C in Trypsin + Collagenase Solution (1xHBSS, 0.05% Gibco^®^ Trypsin+EDTA (Life Technologies™), 20mg/ml collagenase (Sigma)). During that time, they were homogenized by pipetting up and down every 3min. The lysis was stopped by adding 50μl foetal bovine serum and 650μl filter-sterilized Hanks’ solution (1xHBSS, 0.25% BSA, 10mM HEPES pH8). The cells were rinsed with 1ml Hanks’ solution and passed through a 40μm cell strainer (Falcon^®^). They were resuspended in ∼400μl Hanks’ solution with 1:10,000 Hoechst 33258 (Molecular Probes^®^) and transferred to a 5ml polystyrene round bottom tube for FACS sorting. The cells were sorted on FACSAria Fusion sorter by Kevin Clark (MRC WIMM FACS Facility). The gates of GFP (488-530) and DsRed (561-582) channels were set with reference to samples derived from non-transgenic embryos. The fluorescence readouts were compensated when necessary. For ATAC-seq library preparation, the cells were sorted into Hank’s solution. For RNA isolation, the cells were sorted directly into RLT Plus buffer (QIAGEN) + 1% β-mercaptoethanol and processed with the RNEasy^®^ Micro Plus kit (QIAGEN), according to the accompanying protocol. The RNA was quantified and its quality assessed with the use of Agilent RNA 6000 Pico kit. All RNA samples were stored at −80°C.

### SYBR^®^ Green qRT-PCR

3μl of the cDNA diluted in H_2_O were used for technical triplicate qRT-PCR reactions of 20μl containing the Fast SYBR^®^ Green Master Mix (Thermo Fisher Scientific) and appropriate primer pair (see Supplementary Table 1). The reactions were run on 7500 Fast Real-Time PCR System (Applied Biosystems) and the results were analysed with the accompanying software. No-template controls were run on each plate for each primer pair. Each reaction was validated with the melt curve analysis. The baseline values were calculated automatically for each reaction. The threshold values were manually set to be equal for all the reactions run on one plate, within the linear phase of exponential amplification. The relative mRNA levels in each sample were calculated by subtracting the geometric mean of Ct values for housekeeping genes *eef1a1l1* and *ubc* from the average Ct values of the technical triplicates for each gene of interest. This value (ΔCt) was then converted to a ratio relative to the housekeeping genes with the formula 2^-ΔCt^.

### Fluidigm Biomark qRT-PCR

To quantify the differences in *gata2a* expression between wild type and mutant ECs, we crossed homozygous *gata2a*^Δi4/Δi4^ mutants to Tg(*kdrl*:GFP) transgenics to generate Tg(*kdrl*:GFP); *gata2a*^Δi4/Δi4^ embryos. These fish, along with non-mutant Tg(*kdrl*:GFP), were used for FACS-mediated isolation of *kdrl*:GFP^+^ and *kdrl*:GFP^-^ cells to quantitatively compare mRNA expression levels of *gata2a* in the endothelial and non-endothelial cells of wild type and *gata2a*^Δi4/Δi4^ embryos, using the Fluidigm Biomark™ qRT-PCR platform. Briefly, 1ng RNA from FACS-sorted cells was used for Specific Target Amplification in a 10μl reaction with the following reagents: 5μl 2xBuffer and 1.2μl enzyme mix from SuperScript III One-Step Kit (Thermo Fisher Scientific), 0.1μl SUPERase• In™ RNase Inhibitor (Ambion), 1.2μl TE buffer (Invitrogen), 2.5μl 0.2x TaqMan^®^ assay mix (see Supplementary Table 3 for the details of TaqMan^®^ assays). The reaction was incubated for 15min. at 50°C, for 2min. at 95°C and amplified for 20 cycles of 15s at 95°C/4min. at 60°C. The cDNA was diluted 1:5 in TE buffer and stored at −20°C. Diluted cDNA was used for qRT-PCR according to the Fluidigm protocol for Gene Expression with the 48.48 IFC Using Standard TaqMan^®^ Assays (Table S3). Each sample was run in 3-4 biological replicates. The collected data were analysed with Fluidigm Real-Time PCR Analysis software (version 4.1.3). The baseline was automatically corrected using the built-in Linear Baseline Correction. The thresholds were manually adjusted for each gene to fall within the linear phase of exponential amplification, after which they were set to equal values for the housekeeping genes: *rplp0, rpl13a, cops2* ^56^, *lsm12b* ^57^ and *eef1a1l1*. The relative mRNA levels for each sample were calculated by subtracting the geometric mean of Ct values for the housekeeping genes from the Ct value for each gene of interest. This value (ΔCt) was then converted to a ratio relative to the housekeeping genes with the formula 2^-ΔCt^. The ΔCt values were analysed with 2-tailed paired-samples *t*-tests with 95% confidence levels. For Fluidigm Biomark™ qRT-PCR, the ΔCt values were analysed with 2-tailed independent-samples *t*-tests with 95% confidence levels, using IBM^®^ SPSS^®^ Statistics (version 22) software.

### Flow cytometry and isolation of WKM haematopoietic cells

Single cell suspensions of WKM cells were prepared from adult zebrafish kidneys of the required genotypes as described ^58^. Flow cytometry analysis was performed on a FACS Aria II (BD Biosciences) after exclusion of dead cells by uptake of Hoechst dye (Hoechst 33342, H3570, ThermoScientific), as described ^41^. WKM cell counts were performed on a PENTRA ES60 (Hariba Medical) following the manufacturer’s instructions. Note that the cell counter does not recognize the zebrafish nucleated erythrocytes, so these were excluded from this analysis. Cell counts for each genotype were analysed with 2-tailed paired-samples *t*-tests with 95% confidence levels, using a Mann-Whitney test for non-parametric distribution. The scatter plots were generated using GraphPad Prism 8.0 and show medians±SD.

### May-Grunwald and Wright-Giemsa staining

Cell staining with May-Grunwald (MG) stain (Sigma MG500) and Giemsa (GIEMSA STAIN, FLUKA 48900) was performed on haematopoietic cell samples. After cytospin, slides are allowed to air-dry and were stained for 5 min at room temperature with a 1:1 mix of MG:distilled water. Next, slides were drained and stained with a 1:9 dilution of Giemsa:distilled water solution for 30min at room temperature. Excess solution was drained and removed by further washes in distilled water. Finally, the slides were air-dried and mounted in DPX (06522, Sigma) for imaging.

### Whole mount *in situ* hybridization (ISH) and immunohistochemistry

Whole-mount ISH was carried out as described previously ^59^, using probes for *kdrl, runx1, cmyb, gata2a, gata2b, rag1* ^3,36,60,61^, and *gfp* (Supplementary Table 1). For conventional ISH embryos were processed, imaged and the ISH signal quantified as described ^33^. Briefly, the pixel intensity values were assessed for normal distribution with a Q-Q plot and transformed when necessary. Mean values (*μ*) of each experimental group were analysed with 2-tailed independent-samples *t*-tests or with ANOVA with 95% confidence levels, testing for the equality of variances with a Levene’s or Brown-Forsythe test and applying the Welch correction when necessary. For ANOVA, differences between each two groups were assessed with either Tukey’s post-hoc test (for equal variances) or with Games-Howell test (for unequal variances). For all these analyses, the IBM^®^ SPSS^®^ Statistics (version 22) or GraphPad Prism 8.0 package were used. For the analysis of *cmyb* expression in the CHT at 4dpf, the embryos scored as ‘high’ or ‘low’ were tested for equal distribution between morphants and uninjected controls or among wild type, heterozygous and mutant genotypes with contingency Chi-squared tests, applying Continuity Correction for 2×2 tables, using IBM^®^ SPSS^®^ Statistics (version 22).

For fluorescent ISH (FISH) combined with immunohistochemistry, ISH was performed first following the general whole mount *in situ* hybridisation protocol. The signal was developed with SIGMAFAST Fast Red TR/Naphthol, the embryos rinsed in phosphate-buffered saline with tween20 (PBT) and directly processed for immunohistochemistry. Embryos were blocked in blocking buffer (5 % goat serum/0.3 % Triton X-100 in PBT) for 1 hour at RT before incubated with primary antibody against GFP (rabbit, 1:500, Molecular Probes), diluted in blocking buffer overnight at 4°C. Secondary antibody raised in goat coupled to AlexaFluor488 (Invitrogen) was used in 1:500 dilutions for 3h at RT. Hoechst 33342 was used as a nuclear counterstain.

Fluorescent images were taken on a Zeiss LSM880 confocal microscope using 40x or 63x oil immersion objectives. Images were processed using the ZEN software (Zeiss).

### Fluorescence microscopy and counting of *itga2b*-GFP^high^ and *itga2b*-GFP^low^ cells

For widefield fluorescence microscopy, live embryos were anaesthetised with 160 μg/ml MS222 and mounted in 3% methylcellulose and imaged on a AxioLumar V.12 stereomicroscope (Zeiss) equipped with a Zeiss AxioCam MrM. To count *itga2b*-GFP^high^ and *itga2b*-GFP^low^ cells in the CHT, Tg(*itga2b*:GFP;*kdrl*:mCherry); *gata2a*^Δi4/+^ animals were incrossed and grown in E3 medium supplemented with PTU to prevent pigment formation. At 5dpf, the larvae were anaesthetised with MS222 and the tail was cut and fixed for 1h at room temperature in 4% PFA. Next, the tails were mounted on 35 mm glass bottomed dishes (MAtTEK) in 1 % low melt agarose and imaged using a 40x oil objective on an LSM880 confocal microscope (Zeiss). Cells in the CHT region were counted manually on Z-stacks as ‘*itga2b*:GFP^low^’ (HSPCs) or ‘*itga2b*:GFP^high^’ (thrombocytes). Genomic DNA from the heads was extracted and used for genotyping as described above. Cell counts for each genotype were analysed with 2-tailed paired-samples *t*-tests with 95% confidence levels, using a Mann-Whitney test for non-parametric distribution. The graphs were generated using GraphPad Prism 8.0 and show medians±SD.

## Acknowledgements

We thank the staff of the Biomedical Services Units (Oxford, Birmingham and Rotterdam) for fish husbandry. We thank Kevin Clark and Sally-Ann Clarke from the WIMM flow cytometry facility for cell sorting. The flow cytometry facility is supported by the MRC HIU, MRC MHU (MC_UU_12009), NIHR Oxford BRC and John Fell Fund (131/030 and 101/517), the EPA fund (CF182 and CF170), and WIMM Strategic Alliance awards G0902418 and MC_UU_12025. We thank the Wolfson Imaging Centre Oxford imaging. The Wolfson Imaging Centre Oxford is supported by the MRC via the WIMM Strategic Alliance (G0902418), the Molecular Haematology Unit (MC_UU_12009), the Human Immunology Unit (MC_UU_12010), the Wolfson Foundation (grant 18272), and an MRC/BBSRC/EPSRC grant (MR/K015777X/1) to MICA – Nanoscopy Oxford (NanO): Novel Super-resolution Imaging Applied to Biomedical Sciences, Micron (107457/Z/15Z). The facility was supported by WIMM Strategic Alliance awards G0902418 and MC_UU_12025. We thank Fatma Kok and Douglas Vernimmen for critical reading of the manuscript. This research was supported by the British Heart Foundation (BHF Oxford CoRE and BHF IBSR Fellowship FS/13/50/30436 to R.M. and M. K.), by a Wellcome Trust Chromosome and Developmental Biology PhD Scholarship (#WT102345/Z/13/Z. to T.D.) and by the MRC MHU programme number MC_UU_12009/8.

## Data Availability

**All data generated or analysed during this study are included in this published article (and its supplementary information files). Accession codes for ATACseq data will be available before publication**.

## Author contributions

T.D., M.K., C.K., E.P. and R.M. designed the study. T.D., M.K., C.K., J.P-Z., K.G.,C.M. and R.M. performed experiments and analyzed the data. J.P-Z., B.F. and K.G. performed experiments. R.R. performed the bioinformatics analyses, T.D. and R.M. wrote the paper and R.P., E.P. and R.M. edited the paper. R.P., E.P. and R.M. secured funding.

## Declaration of interests

The authors declare no competing interests.

## Notes

#### Summary of Updates

Revised version with added experimental data

