## Supplemental Information for "Deletion of a conserved Gata2 enhancer impairs haemogenic endothelium programming and adult haematopoiesis"

**Supplementary Figures**

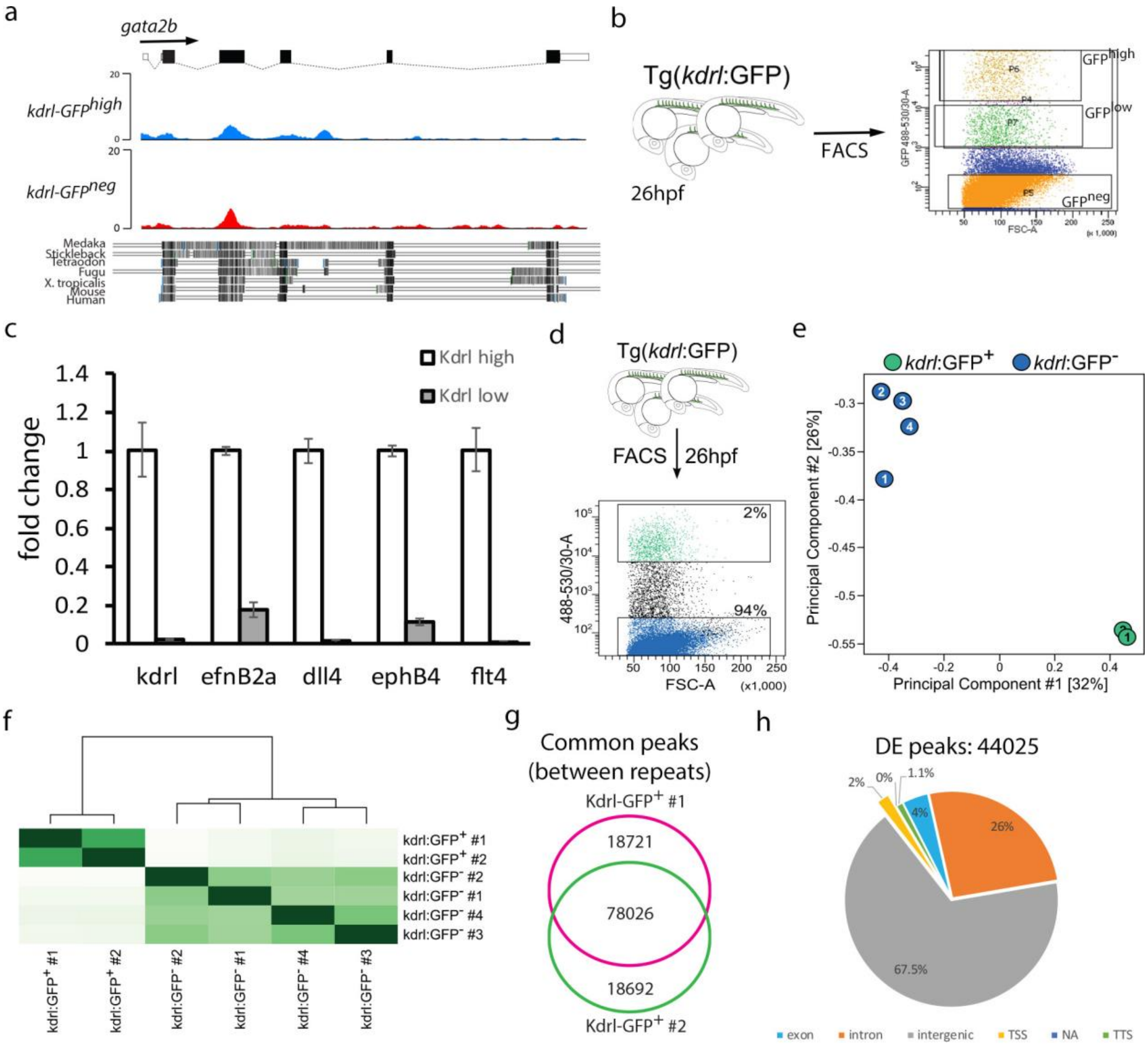


i

Unique EC peaks=5883

| Rank | Motif name | P-value | % of Targets Sequences | Fold enrichment |
|---|---|---|---|---|
| 1 | ERG(ETS) | 1e-1006 | 49.23% | 3.89 |
| 2 | ETS1(ETS) | 1e-779 | 37.57% | 4.21 |
| 3 | ETV1(ETS) | 1e-777 | 42.12% | 3.69 |
| 4 | EWS:ERG-fusion(ETS) | 1e-676 | 30.17% | 4.73 |
| 5 | Fli1(ETS) | 1e-673 | 38.60% | 3.58 |
| 6 | Ets1-distal(ETS) | 1e-580 | 20.53% | 6.46 |
| 7 | GABPA(ETS) | 1e-579 | 30.75% | 3.98 |
| 8 | EWS:FLI1-fusion(ETS) | 1e-563 | 24.87% | 4.91 |
| 9 | EHF(ETS) | 1e-536 | 35.49% | 3.24 |
| 10 | SPDEF(ETS) | 1e-357 | 27.60% | 3.00 |

Common EC and non-ECs peaks =1320

| Rank | Motif name | P-value | % of Target Sequences | Fold enrichment |
|---|---|---|---|---|
| 1 | TR4(NR),DR1 | 1.00E-14 | 4.09% | 3.62 |
| 2 | Sox10(HMG) | 1.00E-11 | 21.44% | 1.50 |
| 3 | PPARE(NR),DR1 | 1.00E-10 | 11.06% | 1.78 |
| 4 | RXR(NR),DR1 | 1.00E-10 | 12.05% | 1.72 |
| 5 | GAGA-repeat | 1.00E-10 | 52.42% | 1.21 |
| 6 | ERG(ETS) | 1.00E-10 | 16.52% | 1.55 |
| 7 | GAGA-repeat | 1.00E-10 | 27.95% | 1.36 |
| 8 | Sox3(HMG) | 1.00E-09 | 21.89% | 1.43 |
| 9 | Nur77(NR) | 1.00E-09 | 3.71% | 2.75 |
| 10 | Erra(NR) | 1.00E-08 | 21.59% | 1.40 |

j

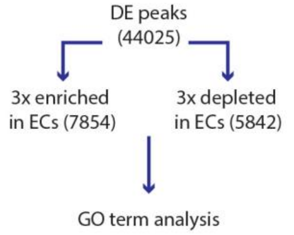


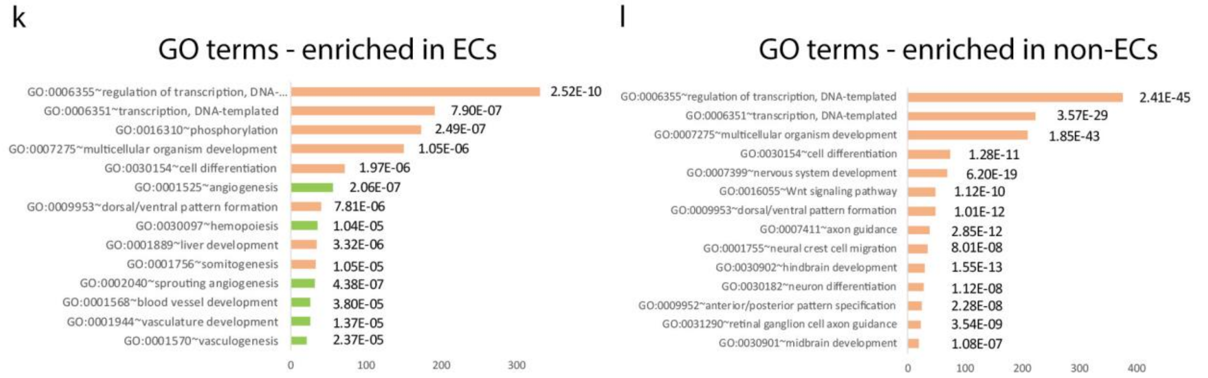

**Supplementary Figure 1. Analysis of open chromatin in endothelial cells by ATAC-seq.** (a) Overview of the zebrafish *gata2b* locus and genome alignment with other species, including mouse and human. Note the conservation within exon sequences in all the species used for comparison. Some intronic sequences are conserved only within fish species. (b) Schematic representation of the gating strategy to test expression of endothelial genes in two distinct *kdrl*:GFP$^{+}$ cell populations: *kdrl*:GFP$^{high}$ (orange, top panel) and Kdrl:GFP$^{low}$ (green). (c) qPCR in kdrl:GFP$^{high}$ and Kdrl:GFP$^{low}$ cell populations, showing significant enrichment for endothelial-specific markers in the kdrl:GFP$^{high}$ population. This population was selected for the ATACseq experiment and will be further referred to as *kdrl*:GFP$^{+}$ for simplicity. (d) *Kdrl*:GFP$^{+}$ (green) and *kdrl*:GFP$^{-}$ (blue) cells were FACS-sorted from 26hpf embryos and used for preparation of ATAC-seq libraries. (e) Principal Component Analysis of 2 *kdrl*:GFP$^{+}$ replicas and 4 *kdrl*:GFP$^{-}$ replicas, showing strong separation of the two cell populations. The same numbers inside the circles denote replicas coming from the same FACS. (f) Clustering of the ATAC-seq samples showing high correlation between *kdrl*:GFP$^{+}$ replicates and among *kdrl*:GFP$^{-}$ replicas. The same numbers represent replicas coming from the same FACS. (g) Venn diagram showing 78026 common peaks between the *kdrl*:GFP$^{+}$ ATACseq profile replicates. (h) Genome-wide distribution of the ATAC peaks from the differential peaks analysis. Note that most peaks are intergenic and intragenic; TSS peaks are only 2% of the total (44026 total peaks). (i) Motif enrichment analysis in endothelial cells (EC-enriched) and common between ECs and non-ECs showing a clear enrichment for ETS binding sites in ECs. (j) Scheme depicting the strategy for the Gene Ontology (GO) term analysis using DAVID [1]. GO term analysis was performed for genes associated to ATACseq peaks that were (k) >3-fold enriched in *kdrl*:GFP$^{+}$ endothelial cells (ECs) or (l) >3-fold depleted in *kdrl*:GFP$^{+}$ cells.

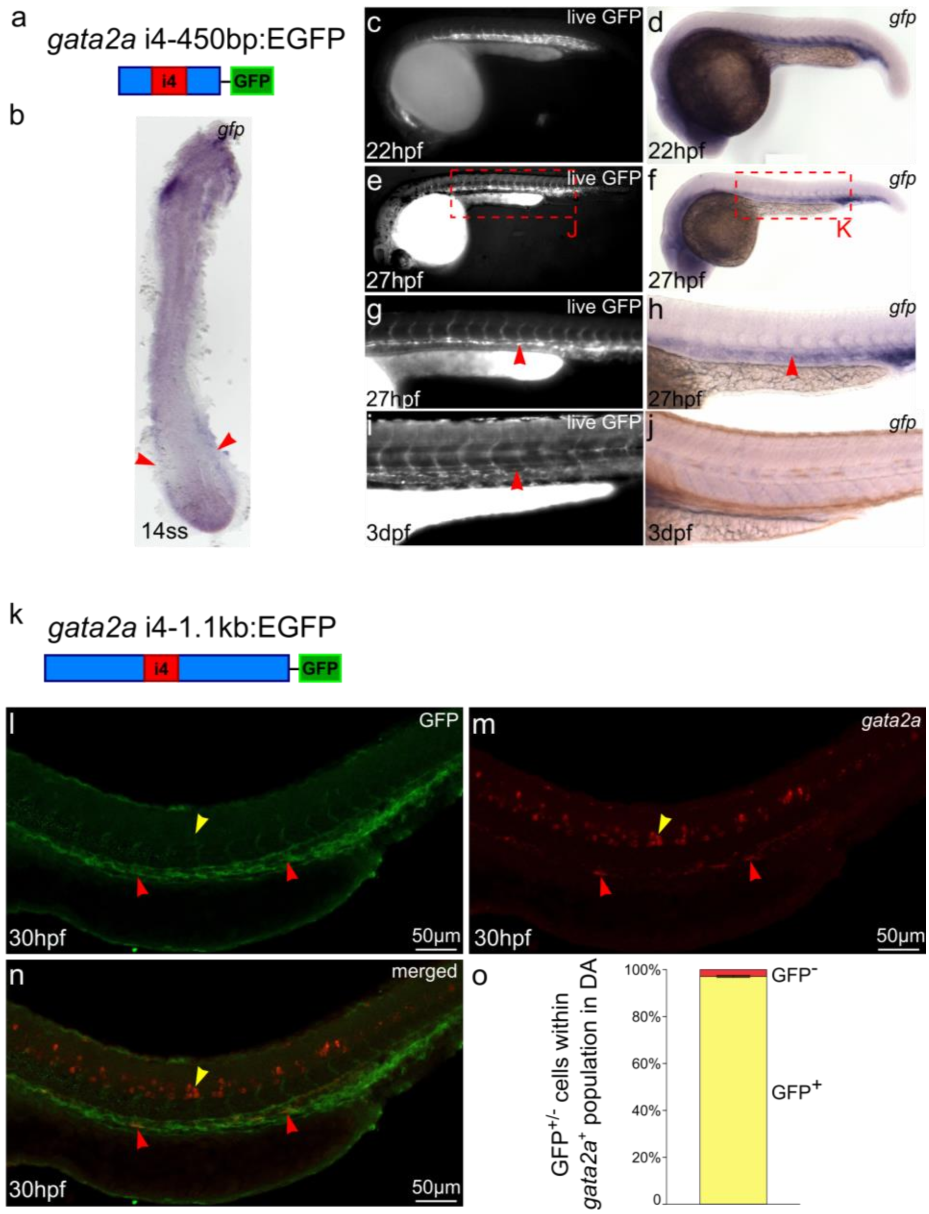


**Supplementary Figure 2. *Gata2a*-i4 enhancer drives GFP expression in PLM, DA and endothelial cells.** (a) Schematic representation of the construct used to generate the Tg(*gata2a*-i4-450bp:GFP) transgenic line. (b) *In situ* hybridization of a 14-somite Tg(*gata2a*-i4-450bp:GFP) embryo (flat mount) shows *gfp* mRNA expression in the posterior lateral plate mesoderm (arrowheads), visible over high background staining in the yolk. (c,d) Fluorescent image of a live Tg(*gata2a*-i4-450bp:GFP) embryo and *in situ* hybridization image of the same embryo probed for *gfp* mRNA at 22hpf reveal activity of the i4 enhancer in the forming dorsal aorta. GFP is also present in the heart. (e-h) At 27hpf, GFP protein was detected in the endothelial cells and is particularly strong in the dorsal aorta (arrowhead in panel g). *In situ* hybridization reveals high degree of overlap between *gfp* RNA and protein expression. (g-h) magnified trunks of embryos in panels e and f as indicated. (i-j) By 3dpf, GFP protein persists in the dorsal aorta (arrowhead) and other endothelial cells, but *gfp* mRNA is undetectable by *in situ* hybridization. (k) Schematic representation of the construct used to generate the Tg(*gata2a*-i4-1.1kb:GFP) transgenic line. (l-n) Confocal images of the trunk of a Tg(*gata2a*-i4-1.1kb:GFP) embryo immunostained with anti-GFP antibody (l) and probed for *gata2a* mRNA (m) at 30hpf, showing an overlap of GFP and *gata2a* in the DA (red arrowheads) but lack of GFP expression in the *gata2a*$^{+}$ neural tube (yellow arrowhead). (n) Merged images from panels l-m. (o) Counting of the *gata2a*$^{+}$ cells represented in panels l-n in 9 embryos shows that >95% of *gata2a*$^{+}$ cells in the DA are also GFP$^{+}$. N=2. Error bars: ±SD.

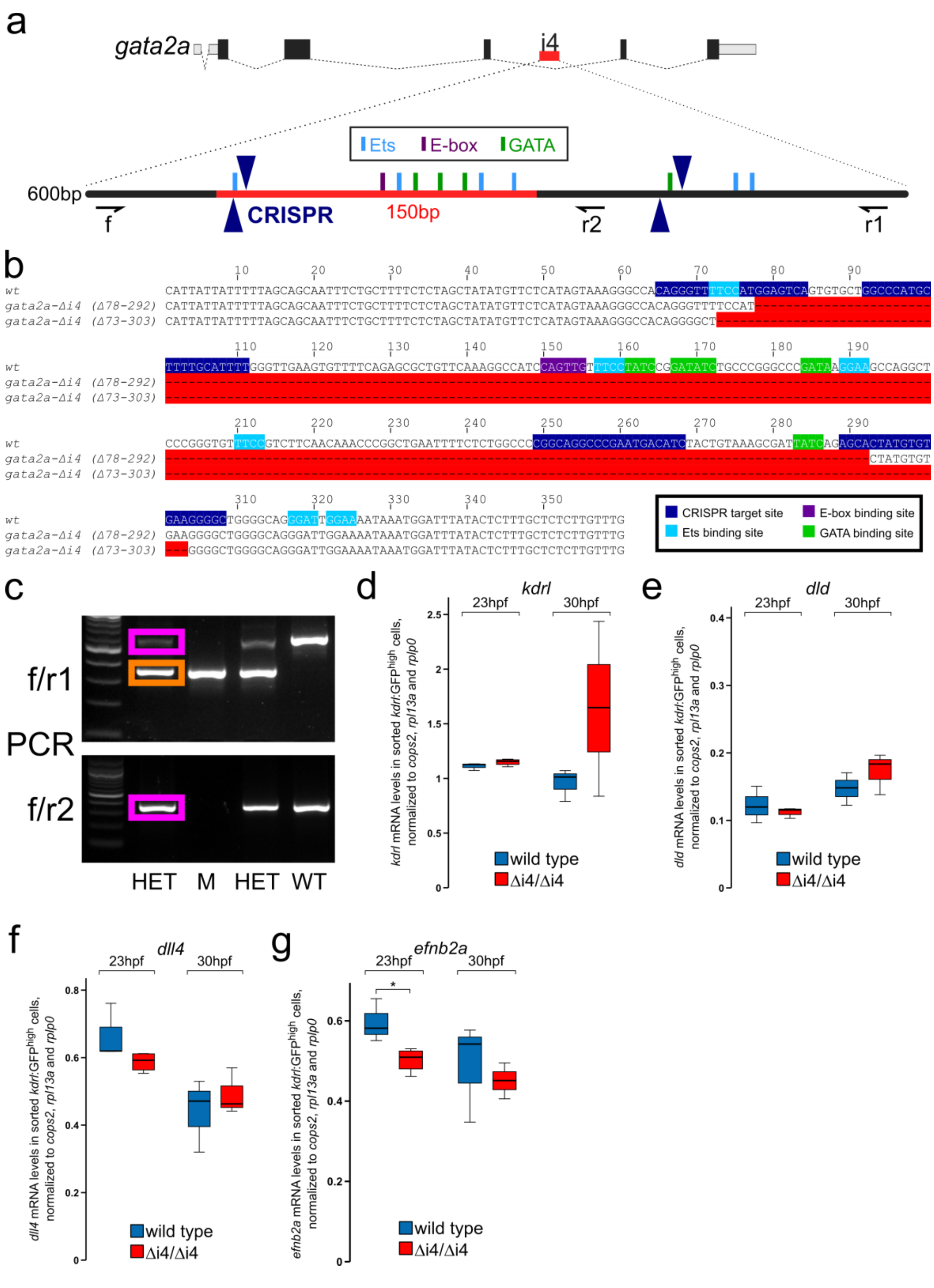


**Supplementary Figure 3. Generation of *gata2a* $^{\Delta i4}$ zebrafish mutants with CRISPR/Cas9 and expression of endothelial markers in *gata2a*$^{\Delta i4/\Delta i4}$ mutants.** (a) Two pairs of CRISPR sgRNAs (dark blue) were designed to flank the highly conserved 150bp region within the *gata2a*-i4 enhancer. Approximate binding positions of diagnostic PCR primers f, r1 and r2 are indicated. Light blue: Ets binding sites; purple: E-box binding sites; green: GATA binding sites (mapped computationally). (b)

Two isolated mutant alleles (Δ78-292 and Δ73-303) carry deletions (red gaps) including the majority of highly conserved transcription factor binding sites. (c) Genotyping of embryos from the incross of two *gata2a*$^{+/\Delta i4}$ mutants with PCR relies on the identification of bands specific to wild type alleles (pink) and the mutant-specific band (orange). Primers f, r1 and r2 are the same as in panel a. The first lane from the left in each gel: 100bp marker. WT: wild type; HET: heterozygote; M: mutant. (d-f) qRT-PCR in sorted *kdrl*:GFP$^{+}$ cells from wt (blue) and *gata2a*$^{\Delta i4/\Delta i4}$ mutants (red) shows no differences in the levels of (d) *kdrl* (23hpf: $t=1.576$, d.f.=5, $p>0.1$; 30hpf: $t=1.399$, d.f.=4, $p>0.2$), (e) *dld* (23hpf: $t=0.585$, d.f.=5, $p>0.5$; 30hpf: $t=1.097$, d.f.=4, $p>0.3$) and (f) *dll4* (23hpf: $t=1.892$, d.f.=5, $p>0.1$; 30hpf: $t=0.734$, d.f.=4, $p>0.5$) in the endothelium of *gata2a*$^{\Delta i4/\Delta i4}$ mutants at 23hpf and 30hpf, compared to wild type. (g) Expression of *efnb2a* was decreased in *gata2a*$^{\Delta i4/\Delta i4}$ mutants at 23hpf ($t=3.008$, d.f.=5, $p<0.05$), but not at 30hpf ($t=0.358$, d.f.=4, $p>0.7$). n=4 for *gata2a*$^{\Delta i4/\Delta i4}$ at 23hpf, n=3 for other samples.

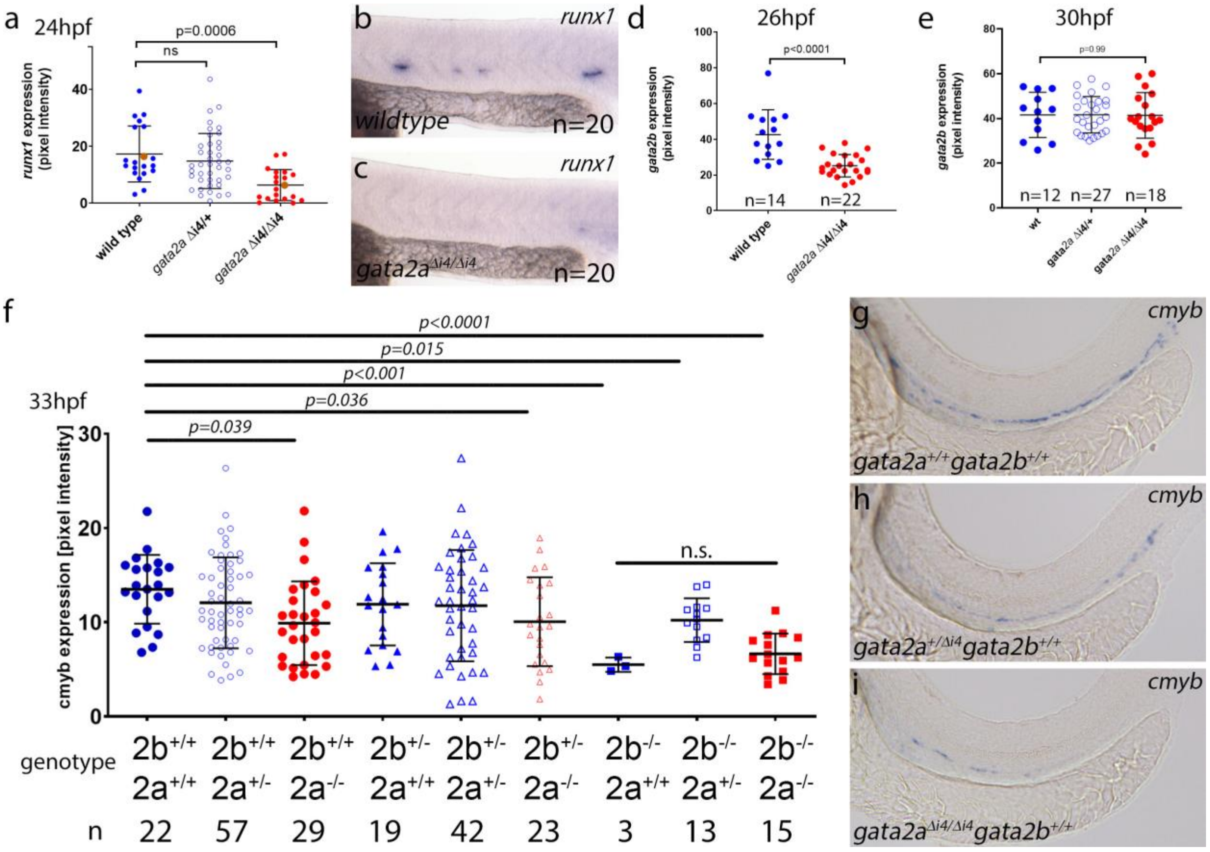


**Supplementary Figure 4. Gene expression analyses in single and double *gata2a*$^{\Delta i4/\Delta i4}$ and gata2b$^{-/-}$ mutants.** (a-c) Analysis of *runx1* expression at 24hpf in *gata2a*$^{\Delta i4/\Delta i4}$ mutants. *Runx1* expression was significantly reduced in *gata2a*$^{\Delta i4/\Delta i4}$ compared to wild type or heterozygous siblings ($\mu_{wt}$=17.3, $\mu_{het}$=14.9, $\mu_{mut}$=6.4; $F$=8.752, d.f.=2, 77; $p$=0.0006; ANOVA). Representative embryos stained for r*unx1* in the (b) wildtype and (c) *gata2a*$^{\Delta i4/\Delta i4}$ mutant embryos (shown in panel a as orange dots). Embryo numbers were n=20, wild type; n=40, *gata2a*$^{\Delta i4/+}$; n=20, *gata2a*$^{\Delta i4/\Delta i4}$. (d) Quantification of *gata2b* expression in wildtype and *gata2a*$^{\Delta i4/\Delta i4}$ mutant embryos at 26hpf. Expression of *gata2b* is significantly reduced in *gata2a*$^{\Delta i4/\Delta i4}$ mutants ($\mu_{wt}$=42.6, $\mu_{mut}$=25.1,; $t$=5.15, d.f.=34; $p$<0.0001; unpaired t test). The number of embryos analysed (n=14, wild type; n=22, *gata2a*$^{\Delta i4/\Delta i4}$) are shown in the panel. (e) Quantification of *gata2b* expression in wildtype and *gata2a*$^{\Delta i4/\Delta i4}$ mutant embryos at 30hpf. We detected no differences in expression of *gata2b* between wild type and *gata2a*$^{\Delta i4/\Delta i4}$ mutants ($\mu_{wt}$=41.56, $\mu_{mut}$=*41*.34,; F=2.54, d.f.=2, 54; $p$=0.99, ANOVA). The number of embryos analysed (n=12, wild type; n=27, *gata2a*$^{\Delta i4/+}$; n=18, *gata2a*$^{\Delta i4/\Delta i4}$) are shown in the panel. (f) Quantification of *cmyb* expression in embryos of all genotypes from *gata2a*$^{\Delta i4/+}$;gata2b$^{+/-}$ incrosses at 33hpf. The expression levels in wildtype, single mutants and double mutant are shown in colour to better highlight the differences. Embryo numbers were n=22, wild type; n=57, *gata2a*$^{\Delta i4/+}$; n=29, *gata2a*$^{\Delta i4/\Delta i4}$; n=19 *gata2b*$^{+/-}$; n=42 *gata2a*$^{\Delta i4/+}$; *gata2b*$^{+/-}$; n=23 *gata2a*$^{\Delta i4/\Delta i4}$; *gata2b*$^{+/-}$; n=3, *gata2b*$^{-/-}$;

n=13 *gata2a*$^{\Delta i4/+}$; *gata2b*$^{-/-}$; n=15 *gata2a*$^{\Delta i4/\Delta i4}$; *gata2b*$^{-/-}$. The data was analysed with a one way Welch's ANOVA test. Note that *cmyb* expression show no statistically significant differences between *gata2b*$^{-/-}$ and *gata2a*$^{\Delta i4/\Delta i4}$; *gata2b*$^{-/-}$ mutants (Dunnett's T3 test for multiple comparisons, t=1.616, d.f.=9.817, p=0.93). Error bars: mean±SD. (g) Representative image of *cmyb* expression in the trunk of 33hpf wildtype, (h) gata2b$^{-/-}$ and (i) *gata2a*$^{\Delta i4/\Delta i4}$; *gata2b*$^{-/-}$ mutants. Pixel intensity of the *in situ* hybridization staining was performed as described [2].

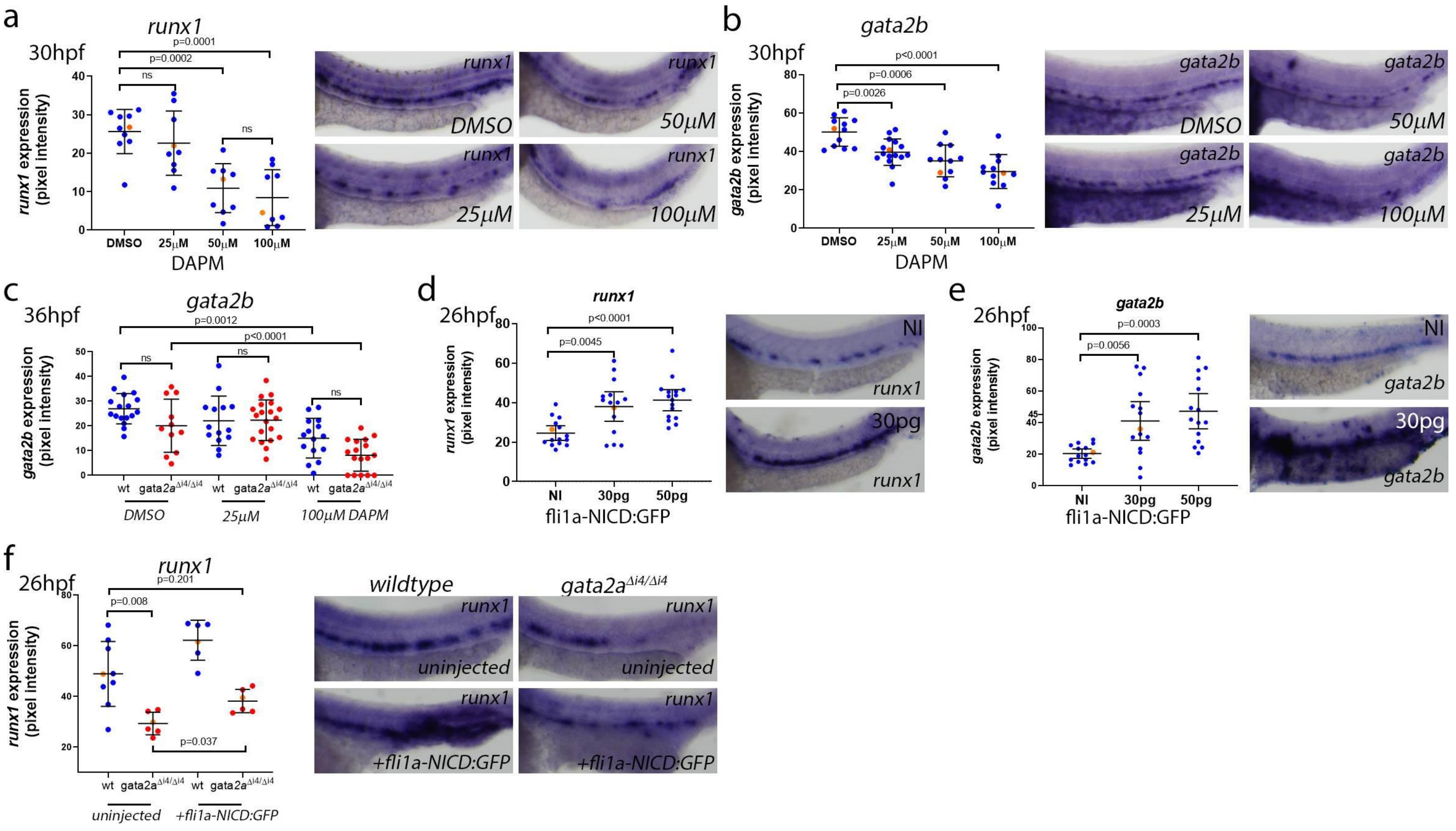
a
runx1
30hpf
p=0.0001
p=0.0002
ns
ns
runx1 expression (pixel intensity)
DMSO
25μM
50μM
100μM
DAPM
b
gata2b
30hpf
p<0.0001
p=0.0006
p=0.0026
gata2b expression (pixel intensity)
DAPM
c
36hpf
gata2b
p=0.0012
p<0.0001
gata2b expression (pixel intensity)
wt
gata2a$^{\Delta i4/\Delta i4}$
DMSO
25μM
100μM DAPM
d
26hpf
runx1
p<0.0001
p=0.0045
runx1 expression (pixel intensity)
NI
30pg
50pg
fli1a-NICD:GFP
e
26hpf
gata2b
p=0.0003
p=0.0056
gata2b expression (pixel intensity)
fli1a-NICD:GFP
f
26hpf
runx1
p=0.201
p=0.008
p=0.037
runx1 expression (pixel intensity)
uninjected
+fli1a-NICD:GFP
wildtype
gata2a$^{\Delta i4/\Delta i4}$

**Supplementary Figure 5. Regulation of *runx1* and *gata2b* in the haemogenic endothelium by Notch signalling.** (a,b) Zebrafish embryos were treated with the Notch inhibitor DAPM from 2-5somite stage until 30hpf at 0 (DMSO control), 25, 50 and 100 μM and expression of *runx1* and *gata2b* analysed by *in situ* hybridization. (a) Analysis of *runx1* expression at 30hpf upon treatment with increasing amounts of the Notch inhibitor DAPM. Expression of *runx1* was significantly reduced upon treatment with 50μM or 100μM DAPM, but not 25 μM DAPM ($\mu_{DMSO}$=25.65, $\mu_{25\mu M}$=22.6, $\mu_{50\mu M}$=10.91, $\mu_{100\mu M}$=8.44; $F$=14.60, d.f.=3, 17.94; $p$<0.0001; Welch's ANOVA). Multiple comparisons performed with a Dunnett's T3 test; significant $p$ values shown in the panel. Embryo numbers were n=10, DMSO; n=9, for the 25μM, 50μM or 100μM DAPM treatment. (b) Analysis of *gata2b* expression at 30hpf upon treatment with increasing amounts of the Notch inhibitor DAPM. *Gata2b* was significantly reduced upon treatment with all the DAPM concentrations tested ($\mu_{DMSO}$=50.13, $\mu_{25\mu M}$=39.62, $\mu_{50\mu M}$=35.09, $\mu_{100\mu M}$=29.54; $F$=13.58, d.f.=3, 24.49; $p$<0.0001; Welch's ANOVA). Multiple comparisons were performed with a Dunnett's T3 test; significant $p$ values are shown in the respective panels. Embryo numbers were n=12, DMSO; n=16, 25μM DAPM; n=11, 50μM DAPM; n=12, 100μM DAPM treatment. (c) Quantification of *gata2b* expression at 36hpf in gata2a$^{\Delta i4/\Delta i4}$ mutants upon DAPM treatment. 100μM DAPM treatment induced a significant decrease in *gata2b* expression in wild type and *gata2a*$^{\Delta i4/\Delta i4}$ mutants. ($\mu_{wt+DMSO}$=26.94, $\mu_{wt+25\mu M}$=22.07, $\mu_{wt+100\mu M}$=15.04; $\mu_{mut+DMSO}$=20.09, $\mu_{mut+25\mu M}$=22.32, $\mu_{mut+100\mu M}$=8.15; Welch's ANOVA). Multiple comparisons were performed with a Dunn's test; significant $p$ values are shown in the respective panels. Embryo numbers were n=17, wt+ DMSO; n=14, wt+25μM DAPM; n=15, wt+100μM DAPM; n=11, *gata2a*$^{\Delta i4/\Delta i4}$ + DMSO; n=20, *gata2a*$^{\Delta i4/\Delta i4}$ +25μM DAPM; n=16, *gata2a*$^{\Delta i4/\Delta i4}$ +100μM DAPM treatment. (d-e) Overexpression of a constitutively active NICD plasmid driven by the endothelial-specific *fli1a* promoter (*fli1a*-NICD:GFP[3]) using 30 and 50pg DNA. (d) *fli1a*-NICD:GFP overexpression increased expression of *runx1* ($\mu_{NI}$=24.51, $\mu_{30pg}$=37.97, $\mu_{50pg}$=41.20; $F$=11.01, d.f.=2, 33.59; $p$<0.0001; Welch's ANOVA, n≥15 for all samples) and (e) *gata2b* in haemogenic endothelium at 30hpf ($\mu_{NI}$=20.42, $\mu_{30pg}$=41.09, $\mu_{50pg}$=47.18; $F$=17.01, d.f.=2, 20.88; $p$<0.0001; Welch's ANOVA, n=15 for all samples). Multiple comparisons were performed with a Dunnett's T3 test; p values are shown in the respective panels. (f) Effects of *fli1a*-NICD:GFP overexpression (30pg DNA) on *runx1* expression between wild type and *gata2a*$^{\Delta i4/\Delta i4}$ mutants at 26hpf. Endothelial-specific NICD overexpression can significantly rescue *runx1* expression in *gata2a*$^{\Delta i4/\Delta i4}$ mutants ($\mu_{wt}$=48.96, $\mu_{mut}$=29.35, $\mu_{wt+NICD}$=62.24, $\mu_{mut+NICD}$=38.21; $F$=25.67, d.f.=3, 12.39; $p$<0.0001; Welch's ANOVA, n≥6 for all samples). Multiple comparisons were performed with a Dunnett's T3 test; $p$ values for each comparison are shown in the respective panels. Error bars: mean±SD; each dot represents the corrected pixel intensity of the *in situ* signal in one embryo. Orange dots correspond to the quantified images shown in each panel.

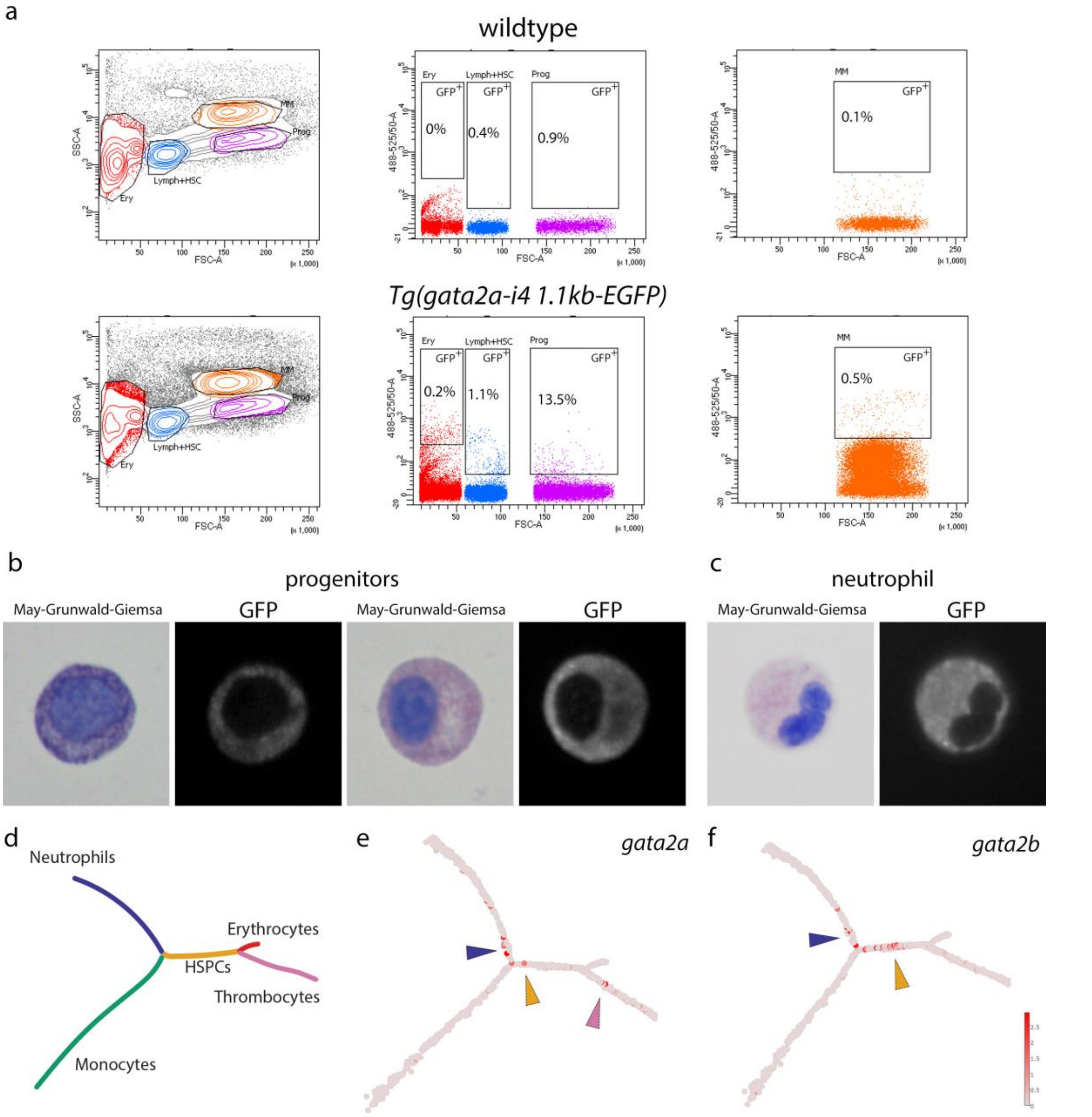


**Supplementary Figure 6. Flow cytometry analysis of GFP expression in Tg(gata2a-i4-1.1Kb:GFP) adult zebrafish.** (a) Plot showing the distribution of cell size (forward scatter) vs granularity (side scatter) to distinguish the various haematopoietic cell populations in the WKM as described [4]. The panels on the right show the gate settings to detect GFP+ cells in the lymphoid gate (Lymph+HSC), progenitor gate (Prog) and myeloid gate (MM). Gata2a-i4-driven expression of GFP was found in all the gates examined. Examples of (b) haematopoietic progenitors and (c) neutrophils expressing GFP, together with May-Grunwald/ Wright-Giemsa staining. (d) Monocle trajectory representing the differentiation path for 5 major cell types in the WKM based on gene expression differences (adapted from [5]). The colours represent each of the branches differentiating from the HSPC 'root'. (e,f) Expression summary for (e) *gata2a* and (f) *gata2b* from single cell RNA-seq data superimposed on the Monocle trajectory. The data is publicly available and was obtained from the BASiCz - Blood

Atlas of Single Cells in zebrafish website (https://www.sanger.ac.uk/science/tools/basicz) [5]. (e) *gata2a* expression was detected in single cells in the neutrophil (blue arrowhead), HSPC (yellow arrowhead) and thrombocyte branch (pink arrowhead). (f) *gata2b* expression was detected in single cells mainly in the neutrophil (blue arrowhead) and HSPC (yellow arrowhead) branches. Expression levels are shown from red to pink (high to low). HSPCs – haematopoietic stem and progenitor cells.

Supplementary Tables

**Supplementary Table 1. Sequences of primers used in this study.**

| Name | Sequence (5'-3') | Purpose | Source |
|---|---|---|---|
| *gata2a*-i4-1.1kb f | GGGGACAAGTTTGTACAAAAAAGCAGGCTCCAGCATCGGGATCCTATAA | Cloning | This study |
| *gata2a*-i4-1.1kb r | GGGGACCACTTTGTACAAGAAAGCTGGGTCACGAATCAAACGCTTTCAG | Cloning | This study |
| *gata2a*-i4-450bp f | GGGGACAAGTTTGTACAAAAAAGCAGGCTGATTGTTGAGAATGTTGTGGTGA | Cloning | This study |
| *gata2a*-i4-450bp r | GGGGACCACTTTGTACAAGAAAGCTGGGTCCAGTCGAGCATGACAAACA | Cloning | This study |
| *gata2a*$^{\Delta i4}$ f | TGGCTAAGTGACCGTCAGAG | Genotyping | This study |
| *gata2a*$^{\Delta i4}$ r1 | TGAAACAAAACGCAGACGAC | Genotyping | This study |
| *gata2a*$^{\Delta i4}$ r2 | GGGTTTGTTGAAGACGGAAA | Genotyping | This study |
| *gfp F* | ACGTAAACGGCCACAAGTTC | Amplify in situ hybridization probe | This study |
| *gfp R* | TGCTCAGGTAGTGGTTGTCG | Amplify in situ hybridization probe | This study |
| *Kdrl F* | CTCCTGTACAGCAAGGAATG | qRT-PCR (SYBR | [6] |
| *Kdrl R* | ATCTTTGGGCACCTTATAGC | qRT-PCR (SYBR) | [6] |
| *efnB2a F* | CCCATTTCCCCCAAAGACTA | qRT-PCR (SYBR) | [7] |
| *efnB2a R* | CTTCCCCATGAGGAGATGC | qRT-PCR (SYBR) | [7] |
| *Dll4 F* | ACGCATACAACCCTAACATGC | qRT-PCR (SYBR) | [8] |
| Dll4 R | CTCTGTCTGCTTCCCACTTTG | qRT-PCR (SYBR) | [8] |
| ephB4 F | CCTGATGAACACGAAAACGGA | qRT-PCR (SYBR) | Bonkhofer et al, unpublished |
| ephB4 R | TGATAGGTCCGCACACTGTT | qRT-PCR (SYBR) | Bonkhofer et al, unpublished |
| Flt4 F | ACAGAGGAGCCATGTTGACA | qRT-PCR (SYBR) | Bonkhofer et al, unpublished |
| Flt4 R | GTCTGGCCTGAGAGTTGAGT | qRT-PCR (SYBR) | Bonkhofer et al, unpublished |
| UbiC F | AAGAGACTCCCATACACCGC | qRT-PCR (SYBR) | Bonkhofer et al, unpublished |
| UbiC R | ATTCTCAATGGTGTCGCTGG | qRT-PCR (SYBR) | Bonkhofer et al, unpublished |
| Ef1a F | GAGAAGTTCGAGAAGGAAGC | qRT-PCR (SYBR) | [9] |
| Ef1a R | CGTAGTATTTGCTGGTCTCG | qRT-PCR (SYBR) | [9] |
| gata2a sgRNA1 | GAAATTAATACGACTCACTATAGGGTGACTCCATGGAAAACCCTGGTTTTAGAGCTA GAAATAGC | Prepare guide RNA template | This study |

| | | | |
|---|---|---|---|
| | | (target region in red) | |
| gata2a sgRNA2 | GAAATTAATACGACTCACTATAGGGGGCCCATGCTTTTGCATTTTGTTTTAGAGCTA GAAATAGC | Prepare guide RNA template (target region in red) | This study |
| gata2a sgRNA3 | GAAATTAATACGACTCACTATAGGGGATGTCATTCGGGCCTGCCGGTTTTAGAGCTA GAAATAGC | Prepare guide RNA template (target region in red) | This study |
| gata2a sgRNA4 | GAAATTAATACGACTCACTATAGGGAGCACTATGTGTGAAGGGGCGTTTTAGAGCTA GAAATAGC | Prepare guide RNA template (target region in red) | This study |
| Universal reverse primer | AAAAGCACCGACTCGGTGCCACTTTTTCAAGTTGATAACGGACTAGCCTTATTTTAAC TTGCTATTTCTAGCTCTAAAAC | Prepare guide RNA template | [10] |
| T7 Gata2b ex3_1 | TAATACGACTCACTATAAATCCGTAGCAACCCGCATCGTTTTAGAGCTAGAAATAGCAAG | Prepare guide RNA template | This study |
| gata2b ex_3 Fwd | CTGTCGATGACGCAACACTG | Genotyping | This study |
| gata2b ex_3 Rev | TGTCGTCATGTTTCCGAGCA | Genotyping | This study |

Most primers were obtained from Sigma-Aldrich. Gata2b primers and guide template were obtained from IDT.

**Supplementary Table 2 –** Open chromatin peaks identified in kdrl-GFP+ (ECs) and kdrl-GFP- (non-ECs) by ATACseq

**Supplementary Table 3. Catalogue numbers of TaqMan® assays used in this work.**

| Gene name | TaqMan® assay catalogue number |
|---|---|
| *cops2* | Dr03114763_m1 |
| *dld* | Dr03111908_m1 |
| *dll4* | Dr03428646_m1 |
| *eef1a1l1* | Dr03432748_m1 |
| *efnb2a* | Dr03073975_m1 |
| *gata2a* | Dr03086718_m1 |
| *gata2b* | Dr03140570_m1 |
| *kdrl* | Dr03432897_m1 |
| *lsm12b* | Dr03139893_m1 |
| *rpl13a* | Dr03101115_g1 |
| *rplp0* | Dr03131546_m1 |
| *runx1* | Dr03074179_m1 |

All TaqMan® assays were obtained from ThermoFisher Scientific.
